# Stimulus-specific plasticity of macaque V1 spike rates and gamma

**DOI:** 10.1101/2020.11.13.381418

**Authors:** Alina Peter, Benjamin J. Stauch, Katharine Shapcott, Kleopatra Kouroupaki, Joscha T. Schmiedt, Liane Klein, Johanna Klon-Lipok, Jarrod R. Dowdall, Marieke L. Schölvinck, Martin Vinck, Wolf Singer, Michael C. Schmid, Pascal Fries

## Abstract

When a visual stimulus is repeated, average neuronal responses typically decrease, yet they might maintain or even increase their impact through increased synchronization. Previous work has found that many repetitions of a grating lead to increasing gamma-band synchronization. Here we show in awake macaque area V1 that both, repetition-related reductions in firing rate and increases in gamma are specific to the repeated stimulus. These effects showed some persistence on the timescale of minutes. Further, gamma increases were specific to the presented stimulus location. Importantly, repetition effects on gamma and on firing rates generalized to natural images. These findings suggest that gamma-band synchronization subserves the adaptive processing of repeated stimulus encounters, both for generating efficient stimulus responses and possibly for memory formation.

## Introduction

Repeated encounters with objects or visual scenes are an everyday experience. As you look around, your eyes often revisit the same visual stimuli on the timescale of seconds (Hooge et al., 2005; Wilming et al., 2013). Thereby, stimulus repetitions are a part of the visual context experienced at any given moment. Prominent theories of visual processing propose that the brain will optimize its responses by making use of the spatiotemporal regularities in the current context (Rao and Ballard, 1999; Schwartz et al., 2007; Snow et al., 2016).

The repetition of an identical stimulus typically leads to reduced firing rates as well as reduced functional MRI signals across numerous brain areas (De Baene and Vogels, 2010; Grill-Spector et al., 2006; McMahon and Olson, 2007; Miller et al., 1993; Sawamura et al., 2005; Solomon and Kohn, 2014; Vogels, 2016; Wissig and Kohn, 2012). These reductions are often referred to as “adaptation”, and they may be indicative of short-term optimization, or alternatively reflect some form of non-beneficial fatigue. They might reflect different plasticity mechanisms, such as simple output fatigue of single neurons, or network changes such as synaptic depression or strengthened inhibitory mechanisms. Notably, some network changes that reduce rates may at the same time strengthen network rhythms. For example, inhibitory mechanisms are tightly linked to gamma-band synchronization (Börgers and Kopell, 2005; Buzsáki et al., 2012). Increased synchronization could maintain or even increase the impact of the reduced number of spikes by increasing their postsynaptic coincidence (Gotts et al., 2012), resulting in an efficient stimulus response with few but effective spikes.

An increase in neuronal gamma-band activity with repeated stimulation has indeed been reported for awake macaque areas V1 and V4 (Brunet et al., 2014). Specifically, local field potential (LFP) gamma-band power increased with the logarithm of the number of repetitions of grating stimuli in V1 and V4, as did V1-V4 coherence and gamma spike-field locking in V4. Yet, key questions remained open. Most importantly, if this phenomenon relates to optimizing stimulus responses, the gamma increase should be specific to the stimulus being repeated, and generalize beyond grating stimuli to initially novel natural images. To address these questions, we recorded LFPs and multi-unit spiking activity (MUA, see Methods) from primary visual cortex (V1) of several macaque monkeys using chronically implanted arrays. We repeatedly presented initially novel, colorful natural images in a pseudorandomly interleaved fashion and found that repetition-related firing rate decreases and gamma-band increases are stimulus-specific. Stimulus specificity was confirmed using uninterrupted sequences of oriented gratings that could change either their stimulus identity or their stimulus position between blocks of 50-100 trials. These paradigms revealed that the stimulus-specific effects had some degree of persistence on the timescale of minutes, and a specificity for the stimulated visual location.

## Results

### Repetition of natural images: task, behavior and average stimulus responses

We investigated stimulus repetition effects using natural and initially novel stimuli, with a paradigm in which different stimuli were interleaved (Fig. 1). Monkeys performed a change detection task on 25 color images of isolated leaves, flowers, sweets, fruits, or vegetables (Fig. S1A-B), which overlapped receptive fields and surrounds (Fig. S1C-E, see Methods for details). For each trial, a stimulus was randomly drawn from a subset of 2-3 stimuli randomly selected from the full set of 25; when a given stimulus had been presented 20 times, it was replaced in the subset by another randomly selected stimulus (see Methods). As not all trials were performed correctly, we used the first 15 correct trials per stimulus for analysis. For a given stimulus, its position in the overall sequence, its neighboring stimuli, and lags between repetitions varied randomly between recording days (Fig. 1B, see Methods), dissociating stimulus-specific repetition effects from potential general effects occurring over the course of a session, and from effects arising from the precise sequence of stimulation.

**Figure 1.**
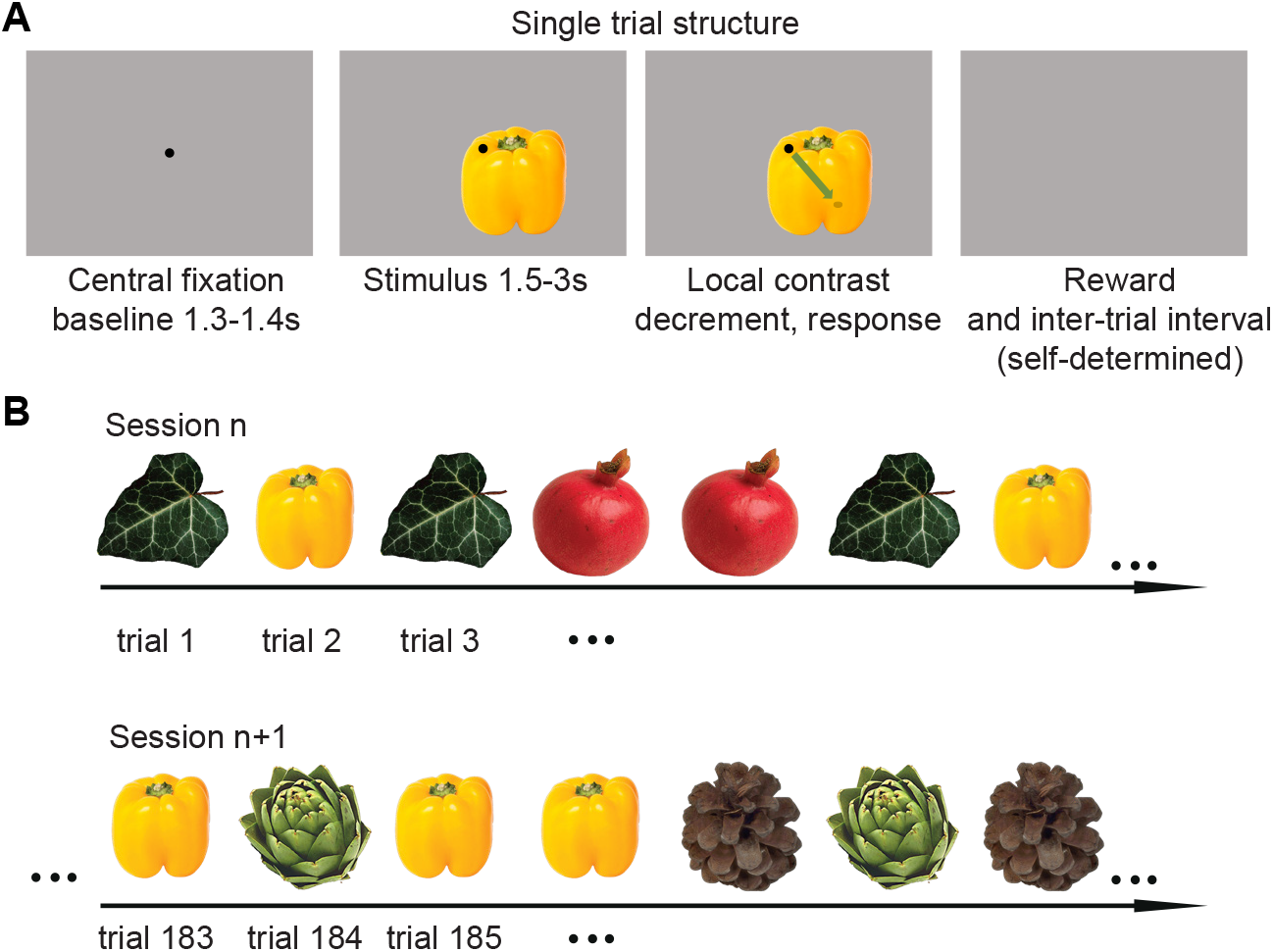
Task for natural image repetition paradigm. (A) Structure of a single trial. Times indicate respective durations. (B) Structure of the trial sequence within and across sessions. Within a session, stimuli could repeat immediately or with up to 4 intervening stimuli. A given stimulus (e.g. the yellow pepper in this example) could occur in different parts of the session on different days, and with different intervening stimuli.

The stimuli were novel to the animals on the first recording day. Although the animals were not required to memorize images, there were clear indications of stimulus memory in the animals’ spontaneous behavior: rapid fixation breaks were frequent during the first few presentations of a novel stimulus, especially on the first day (Fig. S1F-G, detailed analysis in figure legend). This also indicated that monkeys experienced the stimuli as perceptually distinct.

Natural images induced dMUA (for de-noised MUA, see Methods), followed either by sustained responses above the pre-stimulus responses, or by sustained reductions (Fig. 2: examples; Fig. S2E). Both LFP power and MUA-LFP PPC spectra (pairwise phase consistency, see Methods) exhibited clear gamma-band peaks (calculated over 0.5-1.5 s post-stimulus), even for weak MUA responses. LFP and PPC spectra were similar for a given stimulus, and averaging of spectra per stimulus over sites was justified (Fig. S2, Methods), whereas different stimuli varied in gamma amplitude, peak frequency and spectral shape. This variability likely stems from known dependencies of gamma-band activity (“gamma”) on stimulus attributes like spatial frequency, size and structure (Brunet and Fries, 2019; Burns et al., 2011; Gieselmann and Thiele, 2008; Jia et al., 2011; Uran et al., 2020), and in particular on L-M cone contrast (Fig. S2, Peter et al. (2019); Shirhatti and Ray (2018)).

**Figure 2.**
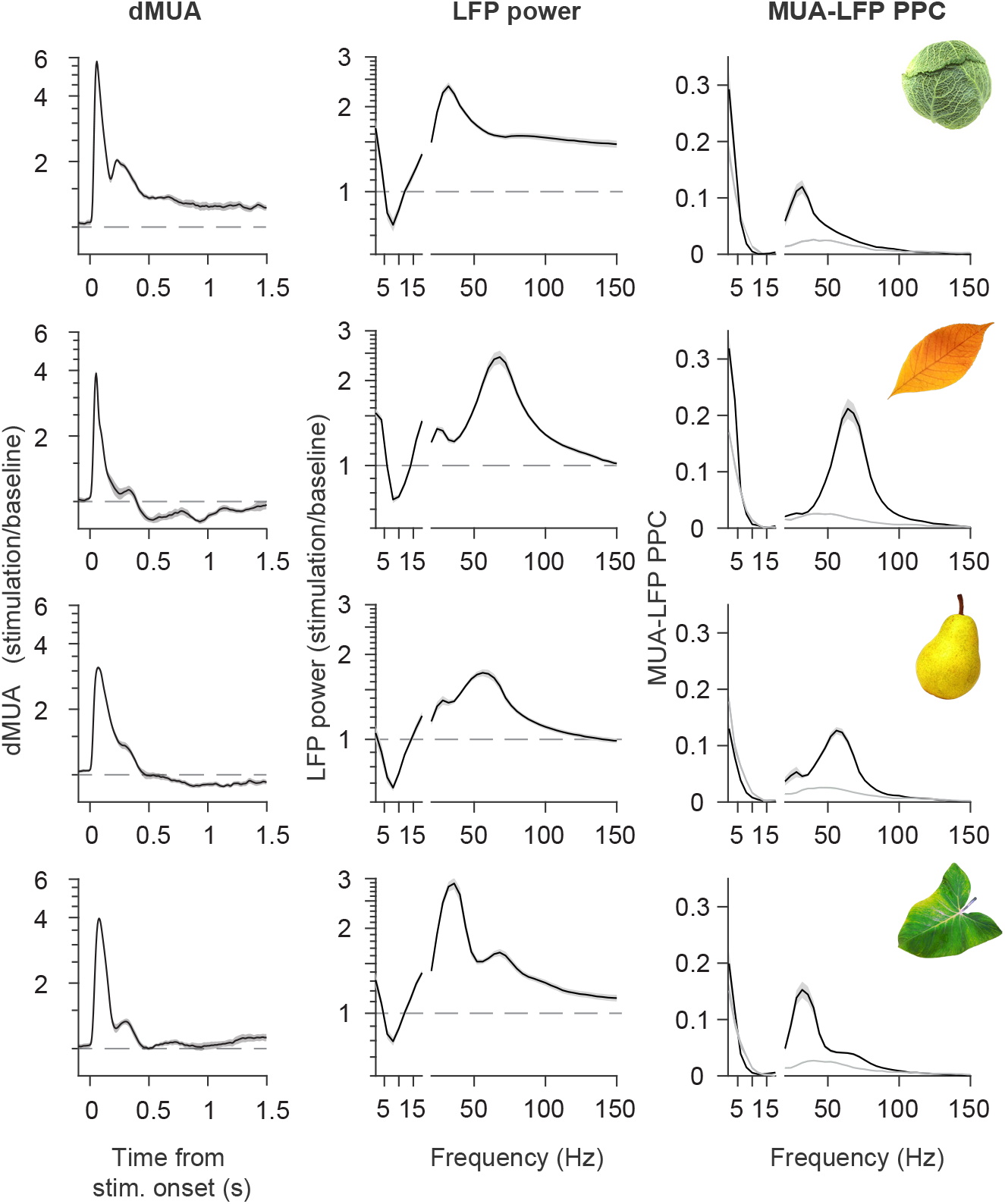
Responses to four example stimuli in monkey H. Example stimuli are shown as insets in the last column. First column: dMUA response. Second column: LFP power change spectra (stimulation/baseline). Third column: MUA-LFP PPC spectra (gray line indicates average baseline activity for the stimulus). Shaded areas indicate ±1 SEM across sessions. Stimulus presentation always lasted from 0 to at least 1.5 s, and spectral analyses used data from 0.5 to 1.5 s.

### Repetition of natural images reduces V1 firing rates

Across recording sites and stimuli, stimulus repetition resulted in decreasing firing rates. The dMUA decrease was particularly strong for the first few repetitions and continued across all analyzed 15 repetitions (Fig. 3A-C; 0.05-1 s post-stimulus onset). To weight all animals, sites and especially stimuli equally, responses were first z-scored and then averaged. As an additional metric of effect size, we also provide the responses normalized by the mean over repetitions (Fig. S3 A-B): firing rates decreased by almost 10% on average. As a new stimulus was introduced at a random time during the session, the strong decrease for the first few repetitions likely indicates stimulus specificity. This will be tested explicitly further below (Fig. 5). Repetition effects were quantified with a simple linear fit, separately for the first four repetitions (“early”) and the later repetitions (“late”), for each stimulus and animal. This separation was motivated by the analysis of gamma power, described further below (see Methods for details). Regression slopes were significantly negative both for early and late repetitions, and more negative for early than late repetitions (Fig. 3B, inset, all p<0.002, two-sided permutation test). The repetition effect survived intervening stimuli: Excluding all immediate repetitions (Fig. 3B, green line) did not change slopes significantly (all p>0.2). The effect had a notably early onset (before 0.1 s post-stimulus onset), and for the early repetitions remained significant throughout the analyzed period (up to 1.5 s post-stimulus, Fig. 3D, E, S3C).

**Figure 3.**
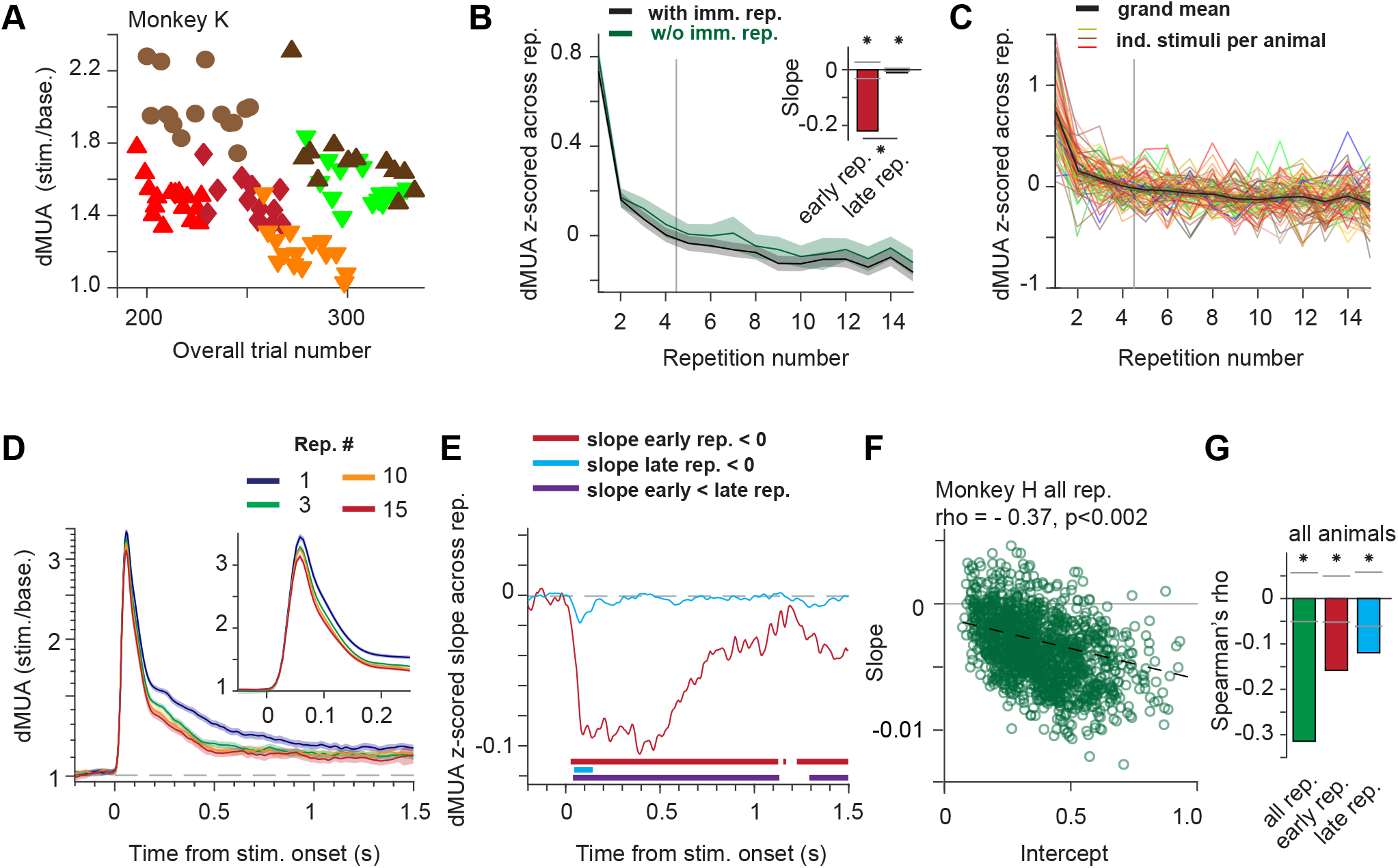
Firing rates during repetition of natural stimuli. (A) Firing rate responses averaged over all sites of monkey K in one example session. Responses to different stimuli are shown with different symbol-color combinations. (B) Black line: Responses as in (A), but averaged over all stimuli, sessions, sites and animals. Green line: Same analysis after removing all immediate repetitions of an identical stimulus. Vertical line separates “early” and “late” repetitions (see main text). Inset: Repetition-related linear slope for early (red) and late (cyan) trials (bars show permutation-based two-sided significance thresholds for p<0.01). (C) Same as black line in (B), with separate lines for each stimulus and animal (line color was chosen to approximate the dominant color in the stimulus). (D) Grand average time-resolved firing rate responses for repetitions 1, 3, 10 and 15. Inset: Zoom on first 250 ms. (E) Repetition-related slopes for early (red) or late (cyan) trials as function of peri-stimulus time. (F) Scatter plot of repetition-related response change (slope) as a function of response strength (intercept), for one example animal. Each circle corresponds to the combination of a recording site with a visual stimulus. Dashed line indicates linear fit. (G) Spearman correlation coefficient between slopes and intercepts, computed as in (F), averaged over animals, for fits to all, early and late trials, respectively. (B-E) All error regions indicate bootstrap estimates of ±2 SEM (see Methods), shown for illustration, whereas statistical inferences were based on non-parametric permutation tests, including correction for multiple comparisons if necessary.

We next asked whether the dMUA repetition effect depended on the response strength of a given dMUA site to a given stimulus. Stronger responses resulted in more pronounced repetition effects regardless of testing all, early or late repetitions separately (based on activity 0.05 to 0.15 s post-stimulus onset, Fig. 3G, 3F for Monkey H, Fig. S3D for Monkeys K and A, all p<0.002). Both response strengths and repetition effects were quantified using linear fits across repetitions: The resulting slopes assess repetition effects, intercepts assess response strengths (cross-validated to avoid regression to the mean, see Methods). To exclude the possibility that the overall correlation derived trivially from poorly responsive site-stimulus combinations, where slope estimates might suffer from a floor effect, we performed a median split by the response intercept, and computed correlations separately for the most and least driven stimulus-site combinations, and separately for slopes fit across all, early or late repetitions (Fig. S3E). The correlation between slopes and intercepts was significantly negative for all combinations, except for the late repetitions in the most driven stimulus-site combinations, which still showed a trend in the same direction. Finally, for some recording sites and stimuli, MUA decreased below pre-stimulus baseline values in the later response period (0.5-1.5 s; Fig. 2, S2). Irrespective of whether the sustained MUA response was above or below the pre-stimulus baseline, the correlation between intercepts and slopes remained significantly negative (Fig. S3F; all p<0.002).

### Repetition of natural images affects V1 gamma-band activity

Repetition effects on gamma-band LFP responses were more varied than effects on firing rates. In particular, for early repetitions, some stimuli induced repetition-related gamma-power decreases, whereas others induced gamma-power increases. For late repetitions, the dominant effect was a gamma-power increase. This is illustrated in Fig. 4A, which shows average LFP power from monkey K for two example stimuli, separately for the indicated repetitions. Repetition-related changes, quantified with a metric (RRC) that normalizes for power changes due to visual stimulation (see Methods), primarily showed increases in the gamma-band, and decreases for higher frequencies likely reflecting spiking activity (Fig. 4B). Gamma-band LFP power, averaged over animals and stimuli after alignment of respective gamma peak frequencies, significantly decreased across early repetitions and then significantly increased across late repetitions (Fig. 4C, responses z-scored as in Fig. 3C for dMUA, all p<0.002 for regression slopes, Fig. S4A; Fig. S4A, responses normalized as in Fig. S3A). Excluding all immediate repetitions (Fig. 4C, green line) did not change slopes significantly (all p>0.14).

**Figure 4.**
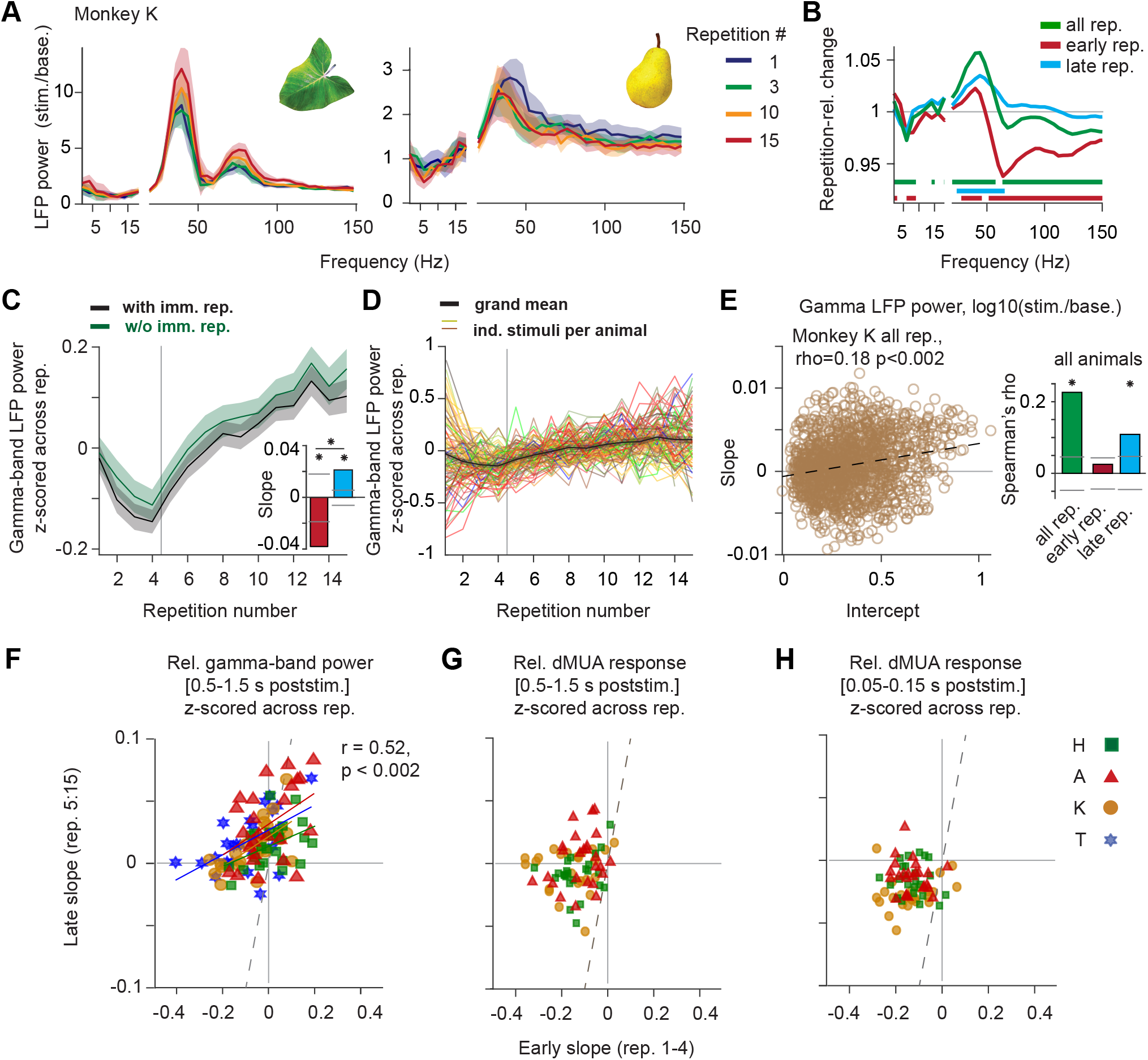
Gamma-band activity during repetition of natural stimuli. (A) LFP power spectra, averaged over all sites and sessions of monkey K, for two example stimuli. Colored lines correspond to the repetition numbers listed on the right. (B) Repetition-related change for early, late or all repetitions (see Methods). (C) Black line: Grand average gamma-band LFP power as a function of stimulus repetition number (see Methods). Green line: Same as black line, but after removing all immediate repetitions of a given stimulus. Dashed line separates “early” from “late” repetitions. Inset: average linear slopes of fits to the early and late repetitions, respectively. (D) Same as black line in (C), with separate lines per stimulus and animal (line colors approximate stimulus color). (E) Scatter plot of repetition-related gamma-power change (slope) as a function of gamma-response strength (intercept), for one example animal. Each circle reflects a stimulus-site combination. Bar plots: Spearman correlation coefficient between slopes and intercepts (computed as in the left panel for monkey K) averaged over animals, for fits to all, early and late trials, respectively. Error regions show ±2 SEM (bootstrap, see Methods), for illustration, whereas statistical inferences were based on non-parametric permutation tests, including correction for multiple comparison if necessary. (F-H) Scatter plots of linear slopes of late versus early trials (see Methods). Each color-symbol combination corresponds to the combination of a stimulus and an animal. Note different axes scales; dashed line is the equality line. (F) Gamma-band LFP power; Colored lines: Significant linear fits per animal. R- and p-values give average over monkeys. (G) dMUA responses for 0.5-1.5 s post stimulus onset; No significant effects. (H) dMUA responses for 0.05-0.15 s post stimulus onset; No significant effects.

Across stimuli, early repetition effects appeared more variable for gamma-band power compared to dMUA (Fig. 4D, Fig. S4B). Given this variability and the reversal of the direction of the repetition effect between early and late trials (Fig. 4C), we investigated the relations between early and late repetition-effect slopes. Early slopes were positively predictive of late slopes (Fig. 4F; r = 0.52, p<0.002, permutation test across stimuli). For dMUA, there was no significant correlation, neither for the time window used for gamma-power analysis (Fig. 4G), nor for the stimulus onset transient period (Fig. 4H; all r < 0.18, all p>0.07). There were also no significant correlations between gamma slopes and dMUA slopes, irrespective of whether the latter were determined for the response onset transient, the gamma-quantification period or the entire trial (all r < 0.21, all p>0.24, corrected for multiple comparisons). Therefore, gamma-power decreases across early trials are unlikely to be explained by the observed co-occuring firing-rate decreases.

Since repetition effects on gamma varied across stimuli, we tested for a relationship between gamma response strength and gamma repetition effects. Stronger responses resulted in significantly more pronounced gamma increases (Fig. 4E, significant for all and late trials, same trend for early trials, analysis as for dMUA in Fig. 3F; Fig. S4C for individual scatter plots). Controls for potential floor-effects left results largely unchanged (Fig. S4D).

We measured the pupil response as an indicator of arousal (Binda et al., 2013; Naber et al., 2013; Peinkhofer et al., 2019). Stimulus-onset induced pupil constriction decreased across early repetitions, with a latency of ~400 ms after stimulus onset (p<0.002, Fig. S4E,F). There were no significant correlations between slopes of dMUA (onset transient, entire trial or LFP time window) and pupil responses, nor between the slopes of gamma-band responses and pupil responses (all r < 0.24, all p ≥ 0.10, one-sided and uncorrected for multiple comparisons).

### Natural image repetition effects in gamma and dMUA are stimulus specific

We have shown that the repetition of natural images induces gamma and dMUA changes which varied across stimuli. That is, for each stimulus, there was a characteristic repetition-related change trajectory. We probed the reliability of these trajectories to explicitly test whether repetition effects were stimulus specific. Stimulus specificity implies that the trajectory for a given stimulus should be relatively reliable, irrespective of when in a session the respective stimulus was presented and with which other stimuli it was interleaved. We calculated split-half correlations between sessions (Fig. 5A, see Methods) of trajectories for LFP power, MUA-LFP PPC and dMUA (z-scored over repetitions to avoid trivial correlations due to offsets between stimuli; permutation test).

**Figure 5.**
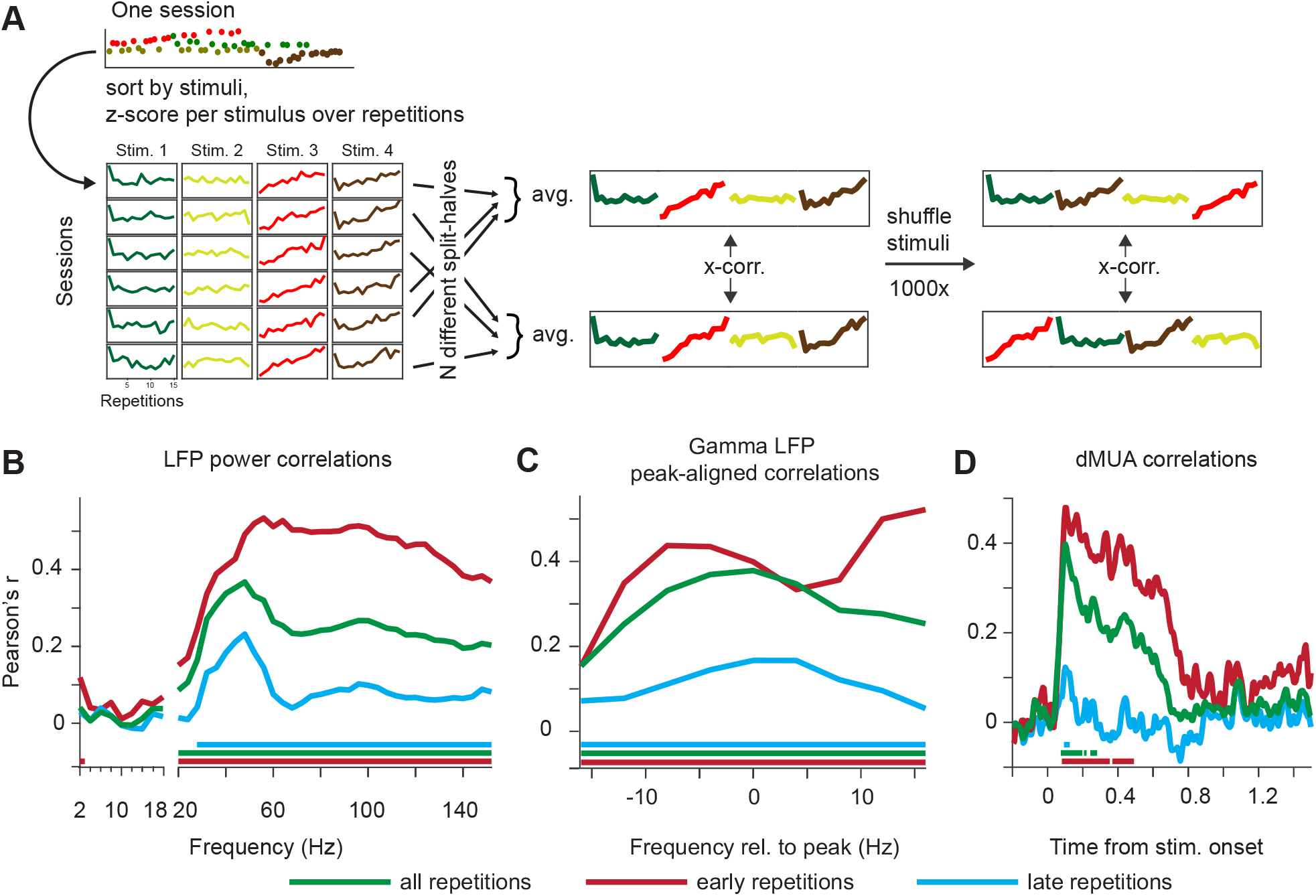
Cross-session correlation of effects of natural stimulus repetition indicates stimulus specificity. (A) Illustration of procedure (see Methods for full details). (B) Correlation spectra for LFP power. (C) Gamma-peak aligned versions of (B). (D) Correlation time courses for dMUA. (B-D): Green lines: all repetitions; Red lines: early (1-4) repetitions; Cyan lines: late (5-15) repetitions. Horizontal bars indicate significance, including multiple comparison correction.

For LFP power, the spectrum of the resulting correlations revealed that stimulus specificity was most pronounced in the gamma band and that it extended into higher frequencies, the latter particularly when including early trials (Fig. 5B); results were confirmed by analysis of spectra aligned to individual stimulus gamma peaks (Fig. 5C). PPC correlation spectra (suggesting stimulus-specific changes in spike synchronization) were more noisy, yet there was a significant correlation in the gamma range when all repetitions were included (Fig. S5A) or when aligning to the individual gamma peak frequencies (for all and for early repetitions, Fig. S5B). For dMUA resolved for time around stimulus onset (Fig. 5D), correlations showed a similar time course as the repetition-related slopes shown in Fig. 3E.

The correlation analyses demonstrate that both MUA and gamma-band responses showed stimulus specificity in their repetition effects. The analysis was optimized to remove variance unrelated to stimulus repetition (by z-scoring across repetitions in a repetition sequence per stimulus, session and site, and by then averaging across session halves). We also developed multiple regression models to fit single-trial responses directly, using either dMUA responses or LFP gamma-band peak aligned responses, averaged across sites, which confirmed the presence of a stimulus-specific repetition effect (all p<0.01, Fig. S5C-E, see legend for details).

### Repetition effects show partial persistence

The results using natural images presented so far showed repetition effects building up despite several intervening stimuli. This suggested partial persistence of effects on the timescale of seconds, and we therefore considered a possible persistence over longer time periods. To maximize sensitivity, we used grating stimuli that are known to induce strong gamma-band responses. Stimuli were presented in a sequence of three blocks of 100 direct stimulus repetitions per block. Between these blocks, switches between two possible stimuli could occur (Fig. 6A,B), without any other changes or breaks in the sequence of trials. Two possible sequences of blocks were presented, ABA or BBA, where A and B signify gratings of different orientation and color (Fig. 6B). Results from specific blocks are referred to using square brackets: e.g., A[B]A denotes the second block in this sequence. Stimuli used as A and B were counterbalanced across sessions, removing effects of stimulus differences in response strength (Fig. 6B). To test for persistence over time, we first document the existence and shape of repetition effects in this paradigm, and demonstrate their stimulus specificity, and then use those properties in the test for persistence.

**Figure 6.**
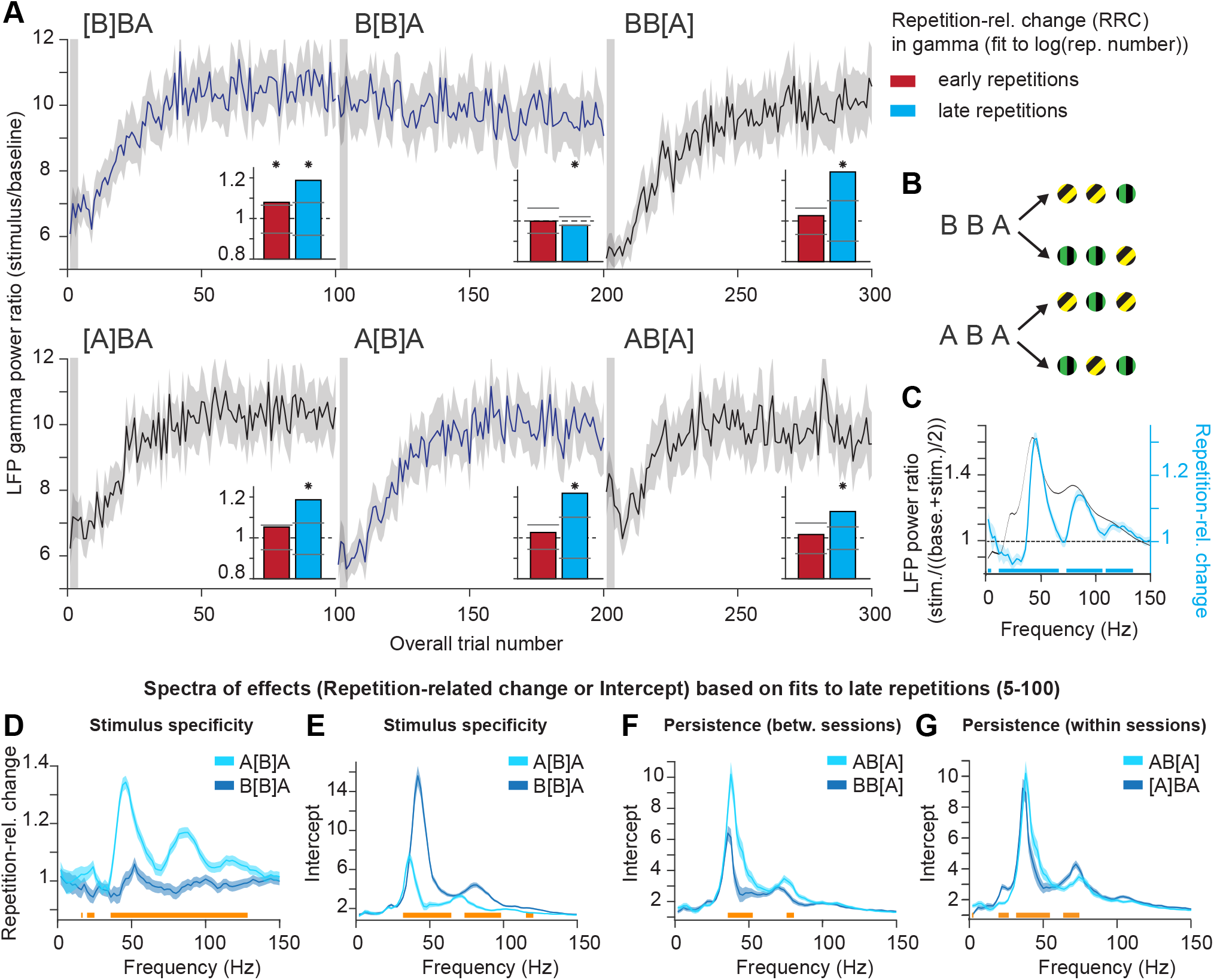
Repetition effects show stimulus specificity and persistence in LFP gamma power. All panels show data averaged over sites, sessions, stimuli and animals. (A) LFP gamma power ratio (stimulation/baseline) as a function of overall trial number. Insets show repetition-related gamma power changes (RRCs, see main text and Methods) for early repetitions (red) and late repetitions (cyan) (bars show permutation-based two-sided significance thresholds for p<0.01). (B) Stimuli used in sequences ABA and BBA. (C) Stimulus-induced LFP power ratio (black) and RRCs (cyan line, horizontal significance bars for test against zero). (D, E) Test for stimulus specificity. (D) Same as repetition-related change spectrum in (C), but for blocks A[B]A, B[B]A. (E) Same as (D) but for intercepts. (F) Test for persistence of effects, using a between-session comparison. Same format as (E), but for blocks AB[A] versus BB[A]. (G) Test for persistence, using a within-session comparison. Same format as (E) but for blocks AB[A] versus [A]BA. (C-G) RRCs and intercepts (the latter estimating initial responses) based on fits to late repetitions. All error regions correspond to ±2 SEM across sessions based on a bootstrap procedure, and are shown for illustration, whereas statistical inferences (horizontal bars below spectra) are based on non-parametric permutation tests (p<0.05), corrected for multiple comparisons.

First, we hypothesized that the first block shows an increase in gamma power with stimulus repetition, and that the rate of this increase becomes smaller with repetition number, replicating previous findings of a log-linear relationship for large numbers of repetitions (Brunet et al., 2014). Correspondingly, further analyses relate neuronal responses to the log-transformed repetition number. Gamma strongly increased in the first block (~50% on average, Fig. 6A). Note that the very first few trials seemed to show no increase or even a decrease, in agreement with the early trials during natural image repetition. For consistency with the natural-image experiments, we define trials 1−4 as “ early trials”, and the remaining trials as “late trials”. All findings reported for late trials also hold when analyzing all trials combined. Within each group of trials, repetition-related changes (RRCs) were quantified as in Fig. 4B (but using log repetition number); this metric normalizes for power changes due to visual stimulation (see Methods). In the first block, for early trials, RRCs in gamma power were significantly positive or trending in this direction (Fig. 6A insets, red bars), and for late trials significantly positive (cyan bars). For the later blocks, for early trials, RRCs showed no significant effects. In block B[B]A, RRCs for late trials were slightly negative. RRCs occurred primarily for gamma and in a similar frequency range as stimulus-induced power changes (Fig. 6C). Gamma peak frequency tended to decrease for early and increase for late trials (Fig. S6A), resulting in shifts in spectral shape also visible in e.g. Fig. 6E.

Second, we hypothesized that repetition-related gamma changes would not transfer to other stimuli, demonstrating stimulus specificity in this paradigm. After a switch to a different stimulus, gamma should again start at a low level (low intercept in regression fit) and increase relatively steeply with repetitions (high slope and therefore RRC). Comparisons of A[B]A and B[B]A showed that gamma-band RRCs were much stronger (Fig. 6D) and intercepts much smaller (Fig. 6E) for A[B]A, keeping stimulus identity, overall trial numbers, time in session, and number of rewards identical.

Third, we hypothesized that a stimulus-specific neuronal assembly that has been repeatedly exposed to a stimulus may show changes that have some “persistence” over minutes (7 min), i.e. that are maintained across many intervening repetitions of another stimulus. That is, a return to a previously shown stimulus should exhibit larger gamma-band intercepts. A respective comparison can be achieved either between sessions or within a session. The between-session comparison, AB[A] versus BB[A], ensures that stimulus identity in the block, overall trial number, time in session, and number of rewards are identical.; the within-session comparison is between AB[A] and [A]BA. Note that the between-session comparison, but not the within-session comparison, controls for the general downward trend for gamma over the course of the session (regression model in Fig. S6E, compare e.g. start of [A]BA to the start of the following bock A[B]A). Both comparisons revealed significant effects: intercepts were larger for AB[A] (Fig. 6F,G), and gamma RRCs were smaller (Fig. S6B,C). Both effects suggest persistence, because changes that persist until a stimulus re-occurs lead to higher initial gamma and require smaller additional changes to reach a plateau. Further, in a single session example, a paradigm with rapid intervening stimulation with several orientations also demonstrated persistence in this case, and a reset of gamma-responses after some minutes of rest (Fig. S6D).

We also tested for stimulus specificity and persistence in gamma-band responses using multiple linear regression modeling (Fig. S6E, see Methods for explanation). The model showed (all p<0.01): 1) a main effect of stimulus repetition, i.e. an increase in gamma-band response with the log-transformed repetition number; 2) stimulus specificity, i.e. an increase in the initial response for the immediate repetition block (B[B]A), and also a net decrease in gamma-band responses for this block for further stimulus repetitions; 3) persistence, i.e. an increase in the initial response for block AB[A], and a reduced increase in gamma for the following repetitions. The model controlled for the effects of overall trial number, pupil responses, microsaccade rates, inter-stimulus-intervals, as well as the stimulus and monkey identity.

Next, we tested for repetition effects, stimulus specificity and persistence in dMUA using the same approaches as for LFP power. Firing rate responses decreased with stimulus repetition (Fig. 7A, B). Repetition-related dMUA reductions were most pronounced for the initial transient, but also significantly affected the response for 1.5 s of stimulation (Fig. 7B). Averaging across the entire stimulation period (0-1.5 s post-stimulus onset) demonstrated that these effects were stimulus specific: RRCs showed a stronger decrease (values below one) and intercepts were higher for A[B]A than B[B]A (Fig. 7C). There was some persistence of this decrease, both between and within sessions (Fig. 7D, E, Fig. S7A, B). These results were confirmed with regression modeling as for gamma (main effects of stimulus specificity and persistence: all p<0.01, Fig. S7C).

**Figure 7.**
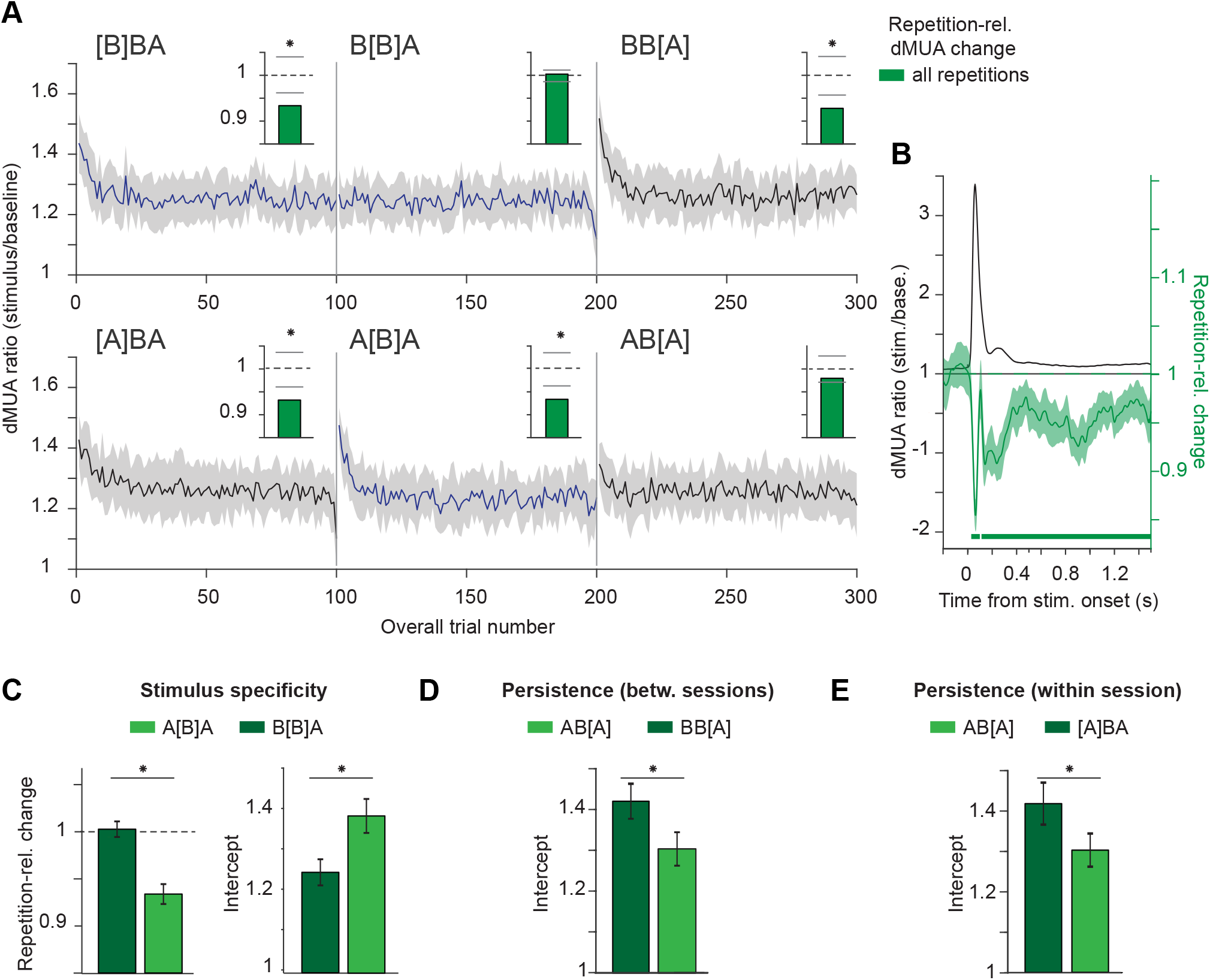
Repetition effects show stimulus specificity and persistence in firing rate responses. All panels show data averaged over sites, sessions, stimuli and animals, all panels except (B) are averaged across the entire stimulation period. (A) dMUA ratio (stimulation/baseline) as a function of overall trial number. Insets show repetition-related dMUA changes across all repetitions (bars show permutation-based two-sided significance thresholds for p<0.01). (B) dMUA ratio (black) and repetition-related changes (green line and horizontal significance bars for test against zero). (C) Test for stimulus specificity based on repetition-related changes and initial responses (i.e. intercepts) (see Methods). (D) Test for persistence, using a between-session comparison. Same format as (C) right panel, but for blocks AB[A], BB[A]. (E) Test for persistence, using a within-session comparison. Same format as (D) but for blocks AB[A], [A]BA. All error regions correspond to ±2 SEM across sessions based on a bootstrap procedure, and are shown for illustration, whereas statistical inferences are based on non-parametric permutation tests (p<0.01, p<0.05 with multiple comparison correction for (B)).

### Repetition effects show location specificity

We hypothesized that repetition-related gamma increases would not transfer to other visual field locations, i.e. that the respective plastic changes were specific to the inducing stimulus location. One grating stimulus was presented in blocks of 50 direct repetitions, either in the RFs of the recorded neurons (In) or outside those RFs (Out). Switches between the two possible locations occurred only between blocks (Fig. 8A-B, further details on block sequence design in the Methods). Specific blocks are denoted by square brackets as in the previous section.

**Figure 8.**
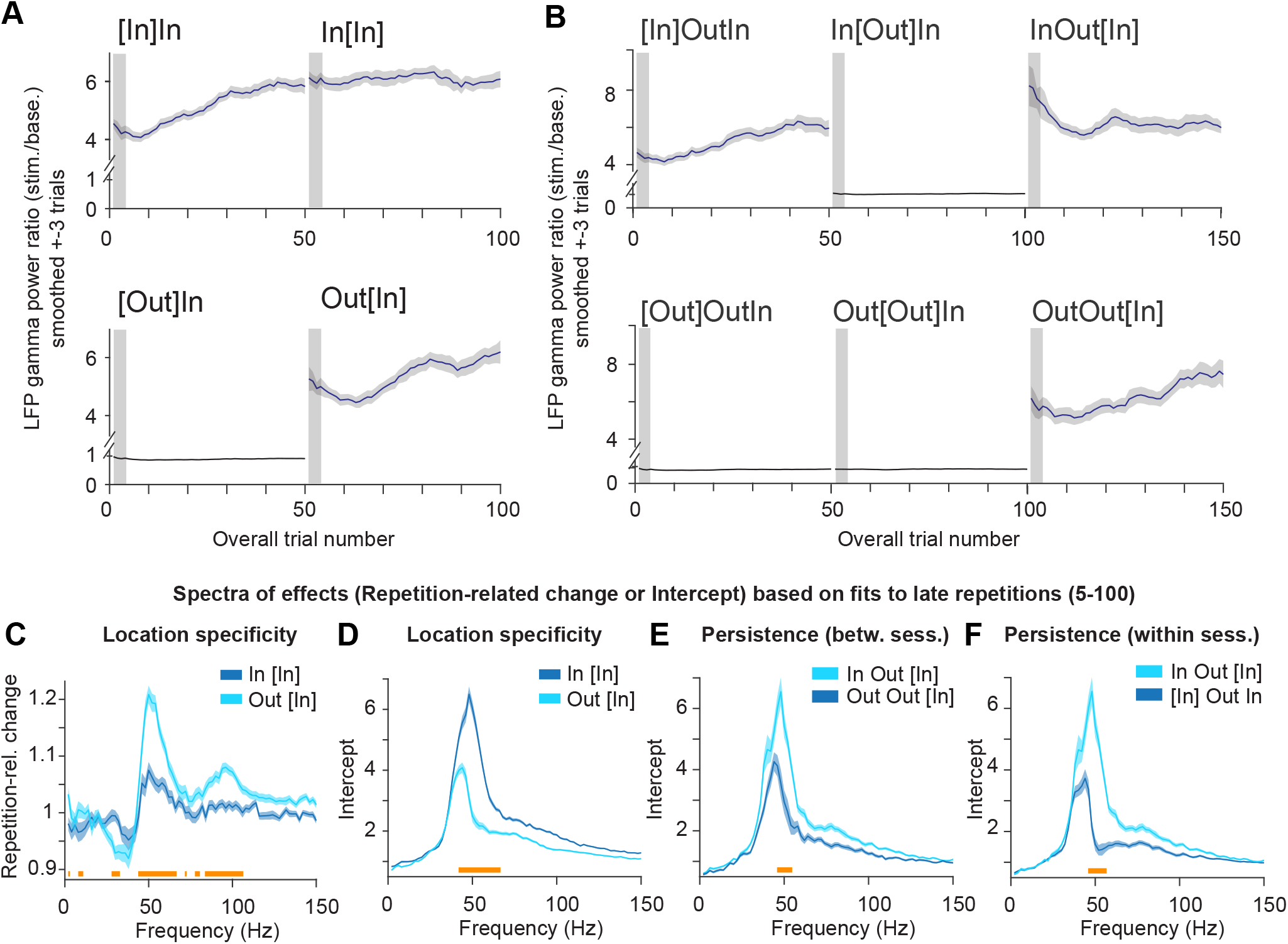
Repetition effects show location specificity and persistence in LFP gamma power. All panels show data averaged over sites, sessions, stimuli and animals, and ±3 trials around the trial number in a given block (see Methods). (A) LFP gamma power ratio (stimulation/baseline) as a function of overall trial number for block sequences InIn and OutIn. (B) Same as (A), but for sequences InOutIn and OutOutIn. (C,D) Similar as Figure 6D, E, but testing for location specificity. (C) Repetition-related change spectrum for late repetitions, for blocks In[In] versus Out[In]. (D) Same as (C) but for the initial response in block (i.e. intercept of the linear regression models). (E, F) Same format as (D), but testing for persistence, using a between-session comparison between blocks InOut[In] and OutOut[In] (E), or a within-session comparison between InOut[In] and [In]OutIn (F). All error regions correspond to ±2 SEM across ±3 trials based on a bootstrap procedure (see Methods), and are shown for illustration, whereas statistical inferences (horizontal bars below spectra) were based on non-parametric permutation tests (p<0.05), including correction for multiple comparisons.

In case of location specificity, after a switch to a different, equi-eccentric location, gamma should again start at a low level (low intercept) and increase relatively steeply with repetitions (high slope). All findings are reported for late trials but also hold when analyzing all trials combined. Contrasting Out[In] versus In[In] showed that this was indeed the case (Fig. 8A, C, D), while controlling for stimulus identity, overall trial number, time in session, and number of rewards.

We tested for persistence of the repetition effect in case of a return to a stimulus location after interrupting *local* stimulation for several minutes (~3.5 min, block InOut[In]). As in the previous section, we tested for persistence within and between sessions, both revealing persistence that bridged one block of 50 trials: intercepts were increased and repetition-related changes decreased, particularly in the gamma band (Fig. 8E, F, S8A, B). Location-specificity and persistence were confirmed using multiple linear regression modeling (all p<0.01, Fig. S8C, see figure legend for details; see Methods).

## Discussion

### Summary

1. Stimulus repetition increased gamma strength and peak frequency, and decreased firing rates in a stimulus-specific manner, suggesting plastic changes predominantly affecting the processing of the repeated stimulus.
2. Repetition-related effects for natural, initially novel stimuli showed stimulus-dependent variation.
3. Repetition effects showed a dependency on initial response strength (a proxy for stimulus drive): whereas stronger initial MUA responses predicted stronger MUA decreases, stronger initial gamma predicted stronger gamma increases.
4. Repetition-related increases in gamma were preceded, for some stimuli, by decreases across the first few repetitions. For the natural image paradigm, most of the dMUA (and pupil response) change occurred with the first repetition.
5. Repetition effects (partially) survived intervening stimuli at the same or a different visual field location, suggesting intermediate-term persistence.
6. Repetition-related changes in gamma were specific to the stimulus location. Also, multi-unit response reductions included the initial response transient. Together, this suggests that the effects originated in early processing stages.

### Repetition-related changes on different timescales

Stimulus repetition might engage different mechanisms with distinct but not mutually exclusive functional benefits, ranging from change or novelty detection to refined processing (Auksztulewicz and Friston, 2016; Friston, 2005; Solomon and Kohn, 2014; Vogels, 2016). Some of these processes should act rapidly, whereas others might build up over time. We observed a pattern of results (points 1 and 4) consistent with the superposition of two processes: a rapid exponential decay of firing rates and gamma, and a slow increase of gamma. For gamma, the relative strength between the fast and slow process might lead to the observed decreases or increases during the first few repetitions, whereas later repetitions may often be dominated by the slow increase. This is supported by the correlation between initial and later repetition-related gamma slopes (Fig. 4F). The superposition of a fast and a slow process of variable strength may explain why previous studies using single natural image repetitions reported (high) gamma decreases (Friese et al., 2012a,b; Kaliukhovich and Vogels, 2012), similar to the average decreases in the first few repetitions of natural images here and novel grating stimuli in the companion paper (Stauch et al., 2020), whereas Brunet et al. (2014) and our Fig. 6 show mostly relatively flat responses to the first few trials of highly familiar grating stimuli followed by increases.

Our study was not designed to investigate the initial, rapid process. We cannot infer whether the strong response to the first presentation of a stimulus reflected an unadapted response or a novelty-driven increase (Kaliukhovich and Vogels, 2014; Vinken et al., 2017; Amado and Kovács, 2016). The concomitant pupil changes point to a potential role of attention or arousal in the natural image paradigm (Binda et al., 2013; Naber et al., 2013; Peinkhofer et al., 2019). For the late repetition effects, the latency of MUA changes, the location specificity, and the regression analyses factoring in microsaccades and pupil response all speak against a purely top-down (attentional) account. Note that repetition-related effects in lower frequencies, rather than gamma, have been linked to task-related changes in top-down processing (Chao et al., 2018; Ghuman et al., 2008; Gilbert et al., 2010; Von Stein et al., 2000; Wang and Dragoi, 2015).

The observed persistence over time scales of seconds to minutes might be related to the number and duration of stimulus presentations. In anesthetized monkey V1, effects of 4 s stimulation on firing rates recovered after 4 s (Patterson et al., 2013). Therefore, the accumulation of repetition effects in the natural image paradigm in awake primate V1, with intervening stimuli and intertrial intervals exceeding stimulus durations, is somewhat surprising; it also exceeds known history effects in rodent and cat V1 (Kim et al., 2019; Nikolic et al., 2009). The observation of partial persistence after many repetitions of the same stimulus may have been facilitated by cumulative effects of many repetitions (Kuravi and Vogels, 2017; Stoelzel et al., 2015). Notably, studies using minute-long stimulus exposure, likely resulting in neuronal fatigue, found stimulus-specific gamma decreases (Jia et al., 2011; Peter et al., 2019). By contrast, studies using short presentations, i.e. one sub-second within-trial repetition of a grating, or several seconds of uninterrupted grating presentation, found gamma increases (Hansen and Dragoi, 2011; Lima et al., 2011; Wang et al., 2011).

Repetition-related increases of rhythmic neuronal activity have also been reported for repetitions of odors: in rodent olfactory and orbitofrontal areas in the gamma-band range (Beshel et al., 2007; Van Wingerden et al., 2010) and in insect olfactory system in the 20 Hz range (Bazhenov et al., 2005; Cassenaer and Laurent, 2007; Laurent et al., 2001; Stopfer and Laurent, 1999). The insect studies found that stimulus repetition leads to a decrease in firing rate and an increase in odor-induced oscillations that is stimulus specific and persistent. While it is unclear whether these effects are related, the prima facie similarity is suggestive of a similar function across species and systems.

### Stimulus dependence of repetition effects

We found that across natural stimuli, stronger initial responses predict stronger repetition-related decreases for firing rates, yet stronger repetition-related increases for gamma. If this relation holds for variability of initial responses that is due to stimulus size, it makes interesting predictions. Increasing the stimulus size of high-contrast stimuli leads to weaker firing rate responses, yet stronger gamma responses, both likely mediated by surround influences (Angelucci et al., 2017; Gieselmann and Thiele, 2008; Jones et al., 2001). Thereby, stimulus size might have opposite effects on repetition-related changes in firing rates and gamma. Large stimuli, inducing relatively weak firing rate responses, should induce relatively weak repetition-related rate decreases. Indeed, the observed firing-rate decreases are relatively small, and previous studies found weak adaptation effects for large, high-contrast stimuli (Wissig and Kohn, 2012; Camp et al., 2009). We also found that the repetition of large stimuli, inducing relatively strong gamma, induced strong repetition-related gamma increases. It will be an interesting question for future studies to systematically investigate the size-dependence of the repetition effects on gamma. As the opposite effects of large, high-contrast stimuli on firing rates and gamma are mediated by surround modulation, a relevant part of the observed repetition-related plasticity might occur in the synaptic mechanisms mediating this surround modulation. Similar considerations might apply to the coloredness of the stimuli. We have previously described that large chromatic, as compared to large achromatic stimuli, induce stronger gamma responses and weaker firing rate responses or even firing rate suppression (Peter et al., 2019).

### Limitations

We did not record single-unit activity. This limits inferences on the cellular origins of increased synchronization. Since different neurons may show different repetition effects, these could average out, potentially masking repetition-related increases or stability in firing rate for subsets of neurons (Homann et al., 2017; Lazar et al., 2018; Patterson et al., 2013; Solomon and Kohn, 2014; Wissig and Kohn, 2012). Notably, Brunet et al. (2014) showed that putative interneurons increasingly synchronized with repetition, whereas poorly driven putative excitatory cells dropped out of the gamma-rhythm. This points to improved coordination of inhibition in stimulus repetition (see below).

### Potential mechanisms of gamma-band increases as adaptive circuit changes

Repetition effects appear to originate from changes at early stages of cortical processing, because 1) the (late) repetition effect is specific to the stimulus and its position, 2) MUA decreases occur with short latency, 3) previous studies found gamma in primates to be generated in V1 rather than LGN (Bastos et al., 2014), and outside the input layers (Xing et al., 2012), 4) subcortical V1 inputs show substantially less, and less stimulus-specific, repetition effects (Sanchez-Vives et al. (2000); Solomon and Kohn (2014), but see Stoelzel et al. (2015)). Since Brunet et al. (2014) also found effects in mid-level area V4, this phenomenon might encompass (or coordinate) several visual areas.

Major candidate mechanisms are output fatigue of single neurons, or network changes (synaptic plasticity) resulting in changes in inhibition and/or excitation. Given 1) the stimulus-specificity of the repetition effect, 2) the small role of lag between repetitions (Fig. 4C), 3) relatively low MUA responses for the natural stimuli, and 4) increased power and peak frequency of gamma, which is itself a network effect, the overall evidence supports a role for network changes (see also Vogels (2016)). What specific form might the network changes take? Stimulus repetition reduces firing rates, but increases synchronization between interneurons (Brunet et al., 2014), as well as gamma power and frequency (this study, Brunet et al. 2014). Frequency shifts likely result from increased input drive (Börgers and Kopell, 2005; Lowet et al., 2017), i.e. stronger input to V1, or synaptic facilitation within V1. Both short-term facilitation and depression have been used to model repetition-related synchronization increases with simultaneous rate decreases (Bazhenov et al., 2005; Wang et al., 2011), in both cases without peak-frequency shifts. Here, we briefly speculate about one possible neuronal mechanism that would result in a process where gamma synchrony is both cause and result of synaptic plasticity (further elaborated in the companion paper Stauch et al. (2020)). A stimulus most strongly drives a subset of excitatory cells in the circuit, which then fire with a short latency. Their drive to interneurons in turn induces a global bout of inhibition that delays (Vinck et al., 2010a) or prevents firing of other cells (De Almeida et al., 2009), setting up a gamma cycle. The most driven cells overcome inhibition first, and therefore repeatedly trigger interneuron firing in subsequent gamma cycles. Thereby, Hebbian spike-timing dependent plasticity strengthens the synapses of strongly driven excitatory neurons onto interneurons (Caporale and Dan, 2008; Hennequin et al., 2017; Huang et al., 2013). Over time, strongly driven cells trigger inhibition more effectively and quickly, increasing gamma power and frequency, and sparsening and synchronizing stimulus responses. Such a process would enhance the impact on downstream areas (Brunet et al., 2014, Stauch et al., 2020) and increase efficiency by reducing overall firing rates. Intriguingly, we note that a distinct class of excitatory neurons combines particularly strong stimulus tuning with strong gamma locking (see Onorato et al. (2020)). The proposal accommodates the drive-dependency observed here across stimuli, and previously across single cells (Brunet et al., 2014), and the partial persistence over minutes of task engagement. The reset of repetition-related changes after a rest period without task engagement (Fig. S6D, Brunet et al., 2014) is in line with the observation that such rest periods may renormalize synaptic changes (Huber et al., 2013; Vyazovskiy et al., 2008). The prevalence of gamma in output layers suggests these as the location of synaptic changes. Other studies have suggested repetition-related strengthening of thalamocortical synapses (Cooke and Bear, 2010; Stoelzel et al., 2015). Such strengthening likely increases drive to cortical neurons and might thereby contribute to the effects observed here. Such effects could be amplified by the proposed cortical mechanisms, which would convert stronger inputs into sparser, more synchronized representations, rather than increasing firing rates.

## ACKNOWLEDGEMENTS

We thank Richard Saunders for planning and performing some of the surgical implants, and Thomas Stieglitz and Eva-Maria Fiedler for producing the polyimide-based ECoG arrays. PF acknowledges grant support by DFG (SPP 1665 FR2557/1-1, FOR 1847 FR2557/2-1, FR2557/5-1-CORNET, FR2557/6-1-NeuroTMR), EU (HEALTH-F2-2008-200728-BrainSynch, FP7-604102-HBP, FP7-600730-Magnetrodes), a European Young Investigator Award, NIH (1U54MH091657-WU-Minn-Consortium-HCP), and LOEWE (NeFF). MCS is supported by DFG Emmy Noether grant SCHM2806/1-1 and ERC starting grant OptoVision 637638, WS is supported by a DFG Koselleck grant and an HFSP collaborative project.

## AUTHOR CONTRIBUTIONS

Conceptualization, A.P., and P.F.; Methodology, A.P., and P.F.; Software, A.P., K.S., J.S., J.R.D.; Formal Analysis, A.P., B.S., M.V., and P.F.; Investigation, A.P., K.S., K.K., L.K., J.K.-L., J.R.D., M.L.S., M. S., W.S., and P.F.; Writing – Original Draft, A.P., and P.F.; Writing – Review & Editing, all authors; Supervision, P.F.; Funding Acquisition, W.S., M.S., and P.F.

## Methods

### Animals

All procedures complied with the German and European law for the protection of animals and were approved by the regional authority (Regierungspräsidium Darmstadt). All animals were group-housed in enriched environments with access to an outdoor space. After the recordings, they continued to live in the facility in their groups. Animal welfare was monitored by veterinarians, technicians and scientists throughout the study. Recordings were made from four male rhesus monkeys weighing 12-16 kg and aged 9-11 years. The recordings used here were from the foveal and parafoveal (up to about 8 degrees visual angle, dva, eccentricity, see Fig. S1C-E) regions of V1 and were made using chronically implanted devices (see below).

### Datasets: tasks and stimuli

Three different datasets were used for this study. Dataset 1 used natural images, was recorded in monkeys K, H, T and A, and is presented in Figures 1–5 and S1-S5. Dataset 2 used many repetitions of gratings to investigate stimulus specificity and persistence, was recorded from monkeys K and H, and is presented in Figures 6–7 and S6-S7. Dataset 3 used many repetitions of gratings to investigate location specificity, was also recorded from monkeys K and H, and is presented in Figures 8 and S8. In all datasets, the monkeys self-initialized trials by acquiring fixation, they performed a change detection task, and correctly performed trials were rewarded with a drop of diluted fruit juice delivered with a solenoid valve system. There were no “catch” trials without changes. The animals were positioned 64-80 cm in front of a 22 inch 120 Hz LCD monitor (Samsung 2233RZ, Wang and Nikolic (2011); Ghodrati et al. (2015)).

#### Dataset 1

Monkeys performed a change detection task on 25 natural images, with repetitions of the images occurring only between trials. Each image was repeated 20 times pseudo-randomly and interleaved with presentations of other repeating images. In the following, the stimuli, and the single-trial, between-trial and between-session design will be described.

##### Stimuli

We chose 25 stimuli from Hemera Photo-Objects Vols. 1, 2, and 3 (Hemera Technologies, see also Woloszyn and Sheinberg (2012)): isolated (cut-out) fruits, vegetables, leaves, flower blossoms and sweets, whose predominant colors were typically red, orange, yellow, green or “dark” (see Figure S1A-B). Stimuli were positioned such that they typically overlapped (slightly) with the fixation point and therefore the fovea (Figure S1C). Images were centered in the lower right quadrant, in accordance with the recorded receptive fields. Stimuli overlapped in a region of about 8 dva diameter, and all available receptive field positions were stimulated.

##### Single-trial structure

On a given trial, a single stimulus was presented after a baseline fixation period of 1.3-1.4 s for a duration of 1.5-3 s (fixation radius 1−1.2 dva), after which a small local change occurred (see Figure 1; parameters that varied between animals are described below). The monkeys could respond to this unpredictable change either by lever release (monkey A) or a saccade to the change location (all other monkeys). Correct responses were followed by a juice reward and the presentation of a grey background screen. The variable time interval between stimulus onset and stimulus change followed a Weibull distribution

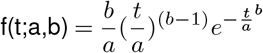

with a = 0.27, b = 2, for positive time points, with a peak probability of a change occurring after around t = 2 s. This resulted in a linear increase in the hazard rate. After this variable duration, a small Gaussian contrast decrement appeared at an unpredictable location on the object. The possible change locations were constrained to positions outside the fixation window and inside the region where all objects overlapped, which was around their center of mass in the lower right visual field and excluded the borders of the object (see Figure S1A-B).

##### Cross-trial task structure

Stimulus repetitions were implemented across trials (rather than within trials). On each day, the same 25 stimuli (described above, see Fig. S1A) were repeated 20 times each in a pseudo-random way, in a sequence with a constrained “lag” (maximally 4 other stimuli between one stimulus and its repetition). This constituted one block of 500 trials. If the monkey worked for more than 500 trials, the task seamlessly proceeded with another stimulus sequence, such that the last images in the block were interleaved with the first images of the second block.

Within a session, the lag between two presentations of the same stimulus was limited, in order to constrain the presentations of a given stimulus to different times of the session on different days (Figure 1B). In order to produce a pseudorandom stimulus sequence with constrained lag, for each trial, a stimulus was randomly drawn from a subset of 2-3 stimuli randomly selected from the full set of 25 stimuli. Once a given stimulus had been presented 20 times, it was replaced in the subset by another stimulus randomly drawn from the full set. Because not all trials were performed correctly, and the analysis was restricted to correct trials, we only analyzed the first 15 correct trials of each stimulus. To dissociate a given stimulus’ first appearance in the session from that of the other images, the following was done. The sequence of each day started with additional, unchanging “dummy” stimuli interleaved with one of the stimuli of the set. The dummies were assigned with a randomly drawn, reduced number of repetitions and therefore would drop out before the completion of the initial stimulus of the set, so that the next stimulus of the set did not start its sequence simultaneously with the first. The images chosen for the second block were produced in an identical manner, with the constraint that they were not identical to the last images in the first block. In cases where this second block was also completed, the last images were interleaved with dummy images.

##### Error handling

Errors were handled such that correct trials, misses, and responses where the target change was initially saccaded to but not held in view, were all counted as repetitions for the purpose of increasing the counter during the task. In all monkeys except monkey T, fixation breaks during the stimulus period resulted in the stimulus turning off immediately to indicate the error. Since monkey T was well-trained on similar tasks, the stimulus was left on the screen for 1 s which allowed the animal to explore the image, which reduced overall fixation break behavior. In all monkeys except monkey T, trials where the monkey fixated correctly for >1 s during the stimulus period were also counted as repetitions. For monkey A, due to limitations in the custom presentation software in the respective laboratory, pseudorandom draws occurred only between two instead of three possible stimuli, and the next two stimuli were introduced after both these stimuli had completed 20 repetitions.

##### Cross-day task structure

To dissociate stimulus-specific repetition effects from potential time-in-session effects, the order in which specific images appeared in the sequence of the day was varied across days (Figure 1B). To generate pseudo-random session sequences of stimuli (i.e. the order of appearance in the session), the first session sequence of images was randomly drawn from the set of images. The following session sequences were drawn pseudo-randomly, with a reduced likelihood of positioning an image at or near positions in the sequence used in previous sequences. This procedure was applied independently for the two blocks. This means that the same image occurred at different points of the session on different days, and had different images shown in neighboring trials on different days. Lag was independent of repetition number and stimulus.

The following parameters varied between animals. For all monkeys except monkey T, stimulus duration ranged from 1.8-3 s, with a peak at 1.9 s, and the fixation point was a blue dot. For monkey T, the timing was 2.3-4 s duration, peak at 2.5 s, the fixation point was a white Gaussian. The task was implemented in custom software for monkey A, MonkeyLogic for monkeys H and K, and an early version of ARCADE for monkey T. The stimulus change was a Gaussian contrast decrement that changed towards the background gray (between 0.8-1.27 dva in size, SD 0.16-0.22) in monkeys A, H, and K. The decrement was smaller and stimulus-dependent in monkey T, based on thresholds from a human observer.

For monkeys T, A, and H, for 1-2 recording days per animal, there were cases of data loss due to technical problems. This left the following number of recording days (blocks) per animal: monkey A 10(10), monkey K 11(20), monkey H 10(19), monkey T 10(19).

#### Dataset 2

Two monkeys performed a change detection task on colored, square-wave grating stimuli (static, spatial frequency 2 cycles per dva, radius 4.5 dva). The stimulus was covering the V1 receptive field locations in the lower right visual quadrant (centered at 4.5/3.5 dva x/y from fixation for monkey H, 3/1 dva x/y for monkey K). In monkey H, the fixation spot was moved up 2 dva from the monitor center, because receptive fields were relatively eccentric, and this allowed the placement of the stimulus on the RFs. Monkeys maintained fixation on a white, circular fixation spot 0.2 dva in size, for 1.3 s of gray background stimulation, followed by 1.5-2.3 s (uniformly distributed) of grating stimulation. In this variable interval, a circular stimulus change of 0.4-0.6 dva diameter (size fixed per session and animal) could occur at a random location on the stimulus. The random locations were restricted such that the full changed spot remained within the stimulus. Monkeys reported the change with a saccade toward the change location within <1 s after the change to obtain a juice reward. The size of the to-be-detected change was kept at a level that resulted in <10% misses. From extended experience with the animals, it was clear that a more challenging task would have led to an increase in the variability of intertrial intervals, potentially interfering with repetition-related effects. The two stimuli analyzed here were chosen to be highly discriminable but to induce similar responses in gamma-band strength and peak frequency. They differed in color and orientation, one green/black grating had a vertical orientation, the other yellow/black grating had an orientation of 60 degrees. Note that these stimuli were not equally bright, nor did they have equal luminance contrast. Grating stimuli were presented in a sequence of three blocks of 100 direct stimulus repetitions per block. Between these blocks, switches between the two possible stimuli could occur (Figure 6). Note that these stimulus switches occurred without any other changes or breaks in the sequence of trials, such that any switch-related changes in neuronal responses could only be due to the switch. Two possible sequences of blocks were analyzed, ABA or BBA, where B signifies a different stimulus from A. Sessions were excluded if they were less than 3 blocks long, or if a break of ≥1 min interrupted stimulation anytime within the 3 blocks. This led to the exclusion of 4 of 48 sessions in monkey K, all due to breaks, and 14 of 62 sessions in monkey H, 6 due to breaks. As a result, Dataset 2 contains 45 sessions of type ABA (monkey H: 23; monkey K: 22) and 47 sessions of type BBA (monkey H: 25; monkey K: 22). In monkey K, a sudden, approximately 5-fold drop in the SNR of MUA was observed after about two weeks of recording for Dataset 2, accompanied by a drop in MUA RF quality. In contrast, clear LFP RFs remained. Therefore, the MUA analyses for this animal (Figure 7) used only 4 sessions of type ABA and 4 sessions of type BBA.

In a single session in monkey H, it was investigated whether the repetition increase would show persistence after an interference block of 25 trials with rapid, repeated stimulation with stimuli of different orientations. To this end, a regular block was followed by an interference block and a second regular block. During the regular block, a vertical, achromatic, maximal contrast, moving square wave grating (1.0 dva spatial frequency, 0.5 dva temporal frequency, 7 dva diameter, position centered on V1 receptive fields, baseline duration 1.3-1.4 s, stimulus duration 1.4 s) was repeated in a passive fixation task for 150 trials. During the interference block, either trials with other orientations (30, 60, 90, 120, 150 degrees angle, first and last interference block in the session) or trials including those other but also the repeated orientation (middle interference block) were shown. Each trial in the interference block consisted of 200 ms individual grating presentations with 100 ms interstimulus interval, such that the total trial duration was equal to trials in the regular blocks. Individual interference trials contained a randomly permuted sequence of orientations. The interference block contained 25 trials, or 125 individual stimulus presentations. After the sequence “regular block, interference block, regular block”, a 10 min break followed, and a new such sequence followed. Figure S6D shows single-trial gamma-band responses using ±10 Hz smoothing (see below for spectral analysis details).

#### Dataset 3

Monkeys H and K performed a change detection task on colored, square-wave grating stimuli (static, spatial frequency 2 cycles per dva, radius 4.25 dva), similar to Dataset 1, but with the following differences. In a given block, one stimulus (with a fixed orientation and color) was shown at one of two different locations: The stimulus was either centered on the V1 receptive field locations in the lower right visual quadrant, or the same stimulus was centered on an equi-eccentric location near the horizontal meridian. The central manipulation of the experiment was that stimulus position could change every 50 correct trials. The design was counterbalanced, such that the overall probability of stimulation occurring at one location or the other was 50%. Sequences of 4 blocks of 50 trials were shown, generating a “task block” of 200 trials. With locations A and B, A being on the V1 receptive fields (In), 8 possible sequences were used in a task block: AAAA, AABB, ABAA, or BAAA (4 types of A-Blocks), and their inverse (all B,… 4 types of B-Blocks). On a given day, A-Blocks and B-blocks alternated, and the starting block alternated between days. The sequence type was assigned in a pseudorandom manner. Blocks of type InIn, OutIn, InOutIn and OutOutIn were analyzed in Figure 8. A ten minute break with a dark monitor followed every 200 trials of stimulation. This encouraged steady responses during task blocks. A further purpose of this break was to use the reset effect of such breaks on gamma-band responses (see also Figure S6D; Brunet et al., 2014). This enabled more data collection on the same recording day. In an additional attempt to ensure independent data in consecutive task blocks, the stimulus was altered on every task block. Specifically, a cyan grating stimulus of 45 degree orientation, a green vertical stimulus, and a yellow 60 degree oriented stimulus was used. Response differences between stimuli are not of interest in this experiment, and all comparisons are made within-stimulus, between-locations and then averaged across stimuli. The stimulus order in the blocks was fixed across days, in order to compare responses to the same stimulus in the same part of the session across days. Task blocks were excluded if they contained fewer blocks than required for analysis, or if a break of ≥1 min interrupted stimulation anytime within these blocks. This led to the exclusion of 3 task blocks, 2 in monkey H and 1 in monkey K. As a result, Dataset 3 contains 30 task blocks of type InIn (monkey H: 18, monkey K: 12), 22 task blocks of type OutIn (monkey H: 12; monkey K: 10), 14 task blocks sessions of type InOutIn (monkey H: 9; monkey K: 5) and 12 task blocks of type OutOutIn (monkey H: 6; monkey K: 6).

### Eye position monitoring

Eye movements from one or two eyes and pupil size (the latter except monkey A) were recorded using infrared illumination. Eye data was recorded with an Eyelink 1000 system (sampling rate 1000 Hz for monkey T, 500 Hz for monkey H and K) or a Thomas Recording system (ET49-B, 122 Hz, monkey A).

### Recordings (electrodes, reference)

For monkeys H and K, recordings were performed with CerePort ("Utah") arrays (64 micro-electrodes; inter-electrode distance 400 μm, tip radius 3-5μ m, impedances 70-800 kΩ, each array containing half of the electrode rows at a length of 1 mm and half at a length of 0.6 mm, Blackrock Microsystems). A reference wire was inserted under the dura toward parietal cortex. For monkey A, a semi-chronic microelectrode array micro-drive was implanted over area V1 of the left hemisphere (SC32-drive from Gray Matter Research; 32 independently movable glass insulated tungsten electrodes with an impedance range of 0.5-2 MΩ and an inter-electrode distance of 1.5 mm, electrodes from Alpha Omega), and the micro-drive chamber was used as the recording reference. For monkey T, we recorded neuronal activity with a micro-machined 252-channel ECoG electrode array implanted subdurally onto areas V1 and V4 of the left hemisphere (252 electrodes; inter-electrode distance 1400μm; electrode diameter 400μm, University of Freiburg) (Rubehn et al., 2009), and we used an electrode adjacent to the lunate sulcus as a recording reference for the section of the array covering area V1 (all analyses are based on local bipolar derivations, see below).

### Recordings (acquisition, filtering)

For monkey A and T, and part of the recordings for monkey K and H, we acquired data with Tucker Davis Technologies (TDT) systems. Data were filtered between 0.35 and 7500 Hz (3 dB filter cutoffs) and digitized at 24,414.0625 Hz (TDT PZ2 preamplifier). For those data, LFP signals were obtained by low-pass filtering and downsampling to 1/24th of the original sampling rate using an 8th order FIR filter. For monkey A, K and H, MUA was obtained by band-pass filtering (300 Hz-12000 kHz) with a 4th order zero-pass Butterworth filter, and filtering and downsampling to 1/24th of the original sampling rate using an 8th order FIR filter. For monkeys H and K in Dataset 1 and parts of Dataset and 3, recordings were acquired using Blackrock Microsystems technology. Channels were amplified, filtered between 0.05 Hz and 10 kHz and digitized at 30 kHz directly at the connector using a CerePlex E headstage (Blackrock Microsystems). Signals were then transferred out of the electrically isolated booth via optic fiber and recorded using a CerebusTM Neural Signal Processor. LFP signals were obtained by low-pass filtering and downsampling to 500 Hz using the Matlab function decimate (8th order zero-phase Chebyshev-filter). MUA was estimated from the broadband signal by band-pass filtering (300 Hz - 30 kHz) with an 8th order zero-phase Chebyshev-filter, rectification, and low-pass filtering and downsampling to 500 Hz using the Matlab function decimate (8th order zero-phase Chebyshev-filter).

For monkeys H, K, and A, the resulting MUA signal is a quasi-continuous measure of high-frequency field power (MUA envelope) and has been used previously by other labs (Legatt et al., 1980; Schmid et al., 2013; Self et al., 2013; Xing et al., 2012).

### Electrode selection and definition of “sites”

To be included in the analysis, channels/sites had to fulfill the following minimal criteria.

1. The site had to have a clear receptive field in the MUA activity, or in the case of monkey T (with the ECoG recordings) in the LFP responses. See section “Receptive field estimation” for a description of the mapping procedures.
2. The RF had to overlap with the presented stimulus. This was ascertained a priori for all experiments by stimulus positioning. In monkey A, this criterion excluded some channels that were not positioned in foveal-parafoveal V1 (lowered below the first encounter of white matter) in an objective manner.

In monkey H, Dataset 2, 4 channels with highly variable SNR, likely due to connectivity issues in the recording system, were excluded. In monkey T, with the ECoG recordings, a few electrodes (9/196) that showed non-physiological responses during the recordings were excluded, and all other electrodes fulfilled the two above-mentioned criteria.

In monkey T (ECoG recordings), all analyses used local bipolar derivations, which are referred to as “(recording) sites”. Sites were only included, if both unipolar electrodes met the above criteria. Local bipolar derivatives were computed between LFPs from immediately neighboring electrodes, as sample-by-sample differences in the time domain, as in previous studies (Bastos et al., 2015; Bosman et al., 2012). If an electrode B had two direct neighbors A and C, it would only be paired with one of these to generate a bipolar site. Thereby, no unipolar recording site entered more than one bipolar site. Additionally, the unipolar sites entering into a bipolar site were required to both originate from the same headstage during recordings. In combination with the above-mentioned exclusion of 9 (unipolar) electrodes, this resulted in 90 recording sites.

For Dataset 1, this resulted in 90 sites in monkey T, 62 unipolar sites in both monkey H and K, and 14 unipolar sites in monkey A. The latter number is relatively low because some of the 32 electrodes were lowered into a part of V1 that covers extremely peripheral regions of the visual field.

For Dataset 2, this resulted in 60 sites in monkey H and 62 sites in monkey K (out of 64 in each case). The same selection was used for Dataset 3.

### Receptive field estimation/Eccentricities

Receptive fields (RFs) were mapped with either bar stimuli (Lima et al. (2010); Peter et al. (2019); monkeys H, K, A), or red dots (monkey T). The signal used for RF mapping was multi-unit activity (MUA) for monkeys H, K, and A, and the LFP gamma power for monkey T. RF eccentricities and positions are displayed in Figure S1C-E.

For monkeys H, K, A (the monkeys with MUA recordings), receptive fields were mapped with moving bar stimuli (spanning the entire monitor). Moving bars (width 1/1/0.1 dva, speed 8/8/17 dva/s, for monkeys K/H/A) were presented in 8 orientations for monkeys H, K and 8-16 orientations for monkey A, each for 10-20 repetitions. MU responses were projected onto the stimulus screen, after shift-correction by the response latency that maximized the back-projected response. MU responses were then fit by a Gaussian function. This Gaussian was used to extract the 10th percentile and the 90th percentile, and this was done separately for each movement direction. Across the 8/16 directions, this yielded 16/32 data points, which were fit with an ellipse. The center of the ellipse was taken as the RF center.

For monkey T (the monkey with ECoG recordings), a red circular stimulus (maximal brightness, RGB [255,0,0], size: 1 dva radius) was presented on a gray background (RGB [128,128,128]) on a grid of positions (with 1 dva steps, i.e. approximately 50% overlap) in the lower right visual field as well as the fovea and the first 1.5 dva above the horizontal meridian (after an initial broader mapping that determined the coverage of the array). Receptive fields were assessed using average relative gamma power from 30-90 Hz (centered approximately around the peak in the spectrum, time window 0.3-4.5 s post-stimulus, power computed with the same parameters as the LFP power described below). To obtain receptive fields, for each channel and each location covered by the grid, relative power was computed by averaging all trials where the stimulus overlapped with the grid location. The receptive field maps of each channel were then normalized by the maximum value, smoothed with a Gaussian (0.25 dva size, SD 0.1 dva), and z-scored. The grid location with the maximal response was taken as the RF center.

### Data analysis

All analyses were done in MATLAB (The MathWorks) using custom scripts and the FieldTrip toolbox (Oostenveld et al., 2011). All randomization or permutation tests were performed with 1000 permutations. All log-transforms have a base of 10.

### Trial selection

Only correctly performed trials were included in the final analyses. In general, repetition number in a sequence could be counted in two ways: 1) counting only correctly performed trials and 2) counting all trials. If incorrectly performed trials have an influence on the repetition effect, the second approach would allocate the repetition to the more accurate position. However, since the most common type of error was a rapid fixation break, this results in missing data in the repetition sequence. We will therefore define repetition here according to the first approach. Exploratory analyses confirmed that approach (2) yields qualitatively similar results. In Dataset 1, we observed that although 20 correct repetitions per stimulus could in theory be performed, substantially less data was present after 15 repetitions according to definition 1) in many cases. We therefore restricted our analyses to the first 15 repetitions.

### Behavioral analysis

Reaction times and correct versus incorrect responses were analyzed using multiple linear regression analyses similar to regression analyses for neuronal data (see below).

### Microsaccade detection and pupil responses

For microsaccade detection, we smoothed horizontal and vertical eye signals (rectangular window of ±5 ms) and differentiated the signals over time points separated by 10 ms to obtain robust eye velocity signals. Data were averaged across eyes. We then used the microsaccade detection algorithm described in (Engbert and Kliegl, 2003) with a velocity threshold of 6*c, where c is the criterion defined as c = Median[v^2^] (Median[v])^2^. Threshold crossings in either the horizontal or vertical direction were considered as microsaccades. We tested several threshold levels and obtained qualitatively similar results.

Pupil size in each trial was computed as the z-score of the pupil size of the single-trial prestimulus baseline (−1 – 0s before stimulus onset) either in a time-resolved manner, for the time period indicated in the figure legend, or, for the regression models, averaged across the entire trial time. Pupil signals across the two eyes were averaged when both eyes were tracked. No pupil data was available for monkey A.

### Spectral analysis (Segmenting Data into Epochs, Calculation of Power and Pairwise Phase Consistency, Peak Estimation)

#### LFP power

For monkeys H and K, activity was re-referenced to the average across the V1 array for LFP power analyses. Unless otherwise noted, all spectra were computed from a fixed part of the trial, namely from 0.5-1.5 s post stimulus onset, or for the baseline (pre-stimulus) period, −1 to 0 s before stimulus onset. Note that the fixation period before stimulus onset had a duration of 1.3-1.4 s, such that the chosen baseline period omits the first 300 ms after fixation, avoiding potential nonlinearities in the response after fixation onset. We excluded the first 500 ms after stimulus onset to minimize effects of transients and non-stationarities on the metrics of rhythmicity and synchronization.

The baseline and stimulus periods were then cut into non-overlapping epochs. Two main types of spectral analyses were performed: 1) Analyses of grating responses for Figures 6–8 and analyses focusing on low-frequency effects (<20 Hz) used 500 ms epochs that were Hann-tapered, and 2) analyses focusing on gamma-band effects in the remaining Figures for natural images, and Figures S6A,D, used 250 ms epochs, and multitaper spectral estimation with 5 tapers taken from the discrete prolate spheroidal sequence, yielding 10 Hz smoothing (Mitra and Pesaran, 1999; Pesaran et al., 2018). Epochs were tapered as described and then Fourier transformed. Power during the stimulation period was normalized to the pre-stimulus baseline period, separately for each site, in the following manner: Power per frequency and per trial was calculated as described above. Power calculated for the pre-stimulus baseline period was then averaged across trials. Finally, trial-wise normalized power was calculated for the stimulation period dividing by the average pre-stimulus spectrum.

#### MUA-LFP phase locking

For MUA-LFP phase locking, only electrodes selected by the procedure described above were used. In addition, for MUA-LFP pairs, we required that the electrodes were direct neighbors in the array. MUA-LFP phase locking was computed as follows. The cross-spectral density between LFP and MUA signal for each trial (cross-spectra) was computed using the same spectral estimation parameters as for the LFP power spectra described above. The cross-spectrum per trial was then normalized by its absolute values, resulting in cross-spectral phases (without amplitude information). Normalized cross-spectra were then used to compute the Pairwise Phase Consistency (PPC), using FieldTrip (Oostenveld et al., 2011). The PPC is unbiased by the trial count (Vinck et al., 2010b). For a given MUA site, the PPC values were then averaged across all the combinations with LFPs from the other selected electrodes that neighbored the respective MUA site. MUA-LFP combinations from the same electrode were excluded to avoid artifactual coherence due to bleed-in of spikes into the LFP (Buzsáki et al., 2012; Ray and Maunsell, 2011). Because of the distance between electrodes (at least 400μ m), this was not an issue for MUA-LFP combinations from different electrodes. Single-trial PPC values were computed across non-overlapping epochs.

#### Determining gamma peak per stimulus

We determined individual gamma peaks per stimulus, because spectra were highly stimulus-specific, with strong variations in gamma-band peak frequency and spectral shape, in particular for natural images (Dataset 1, see Figure 2 and S2). In addition, many spectra featured several distinct peaks for a single image and site. Frequently, these were also visible in MUA-LFP PPC spectra (see Figure 2).

For each stimulus, the largest peak in the gamma-band response was determined for relative power spectra (and with an identical procedure, for the MUA-LFP PPC during the stimulus presentation time). Given that the peak frequency varied between stimuli, this could be used to determine a peak-centered gamma-band for each stimulus in which to average gamma-band activity. Local maxima were determined between 20 and 190 Hz. A peak was defined as a local maximum as implemented in the Matlab function “findpeaks”.

For Dataset 2 and 3 (persistence and location specificity tests), we observed that per Dataset, stimulus and animal, >99% of site-session combinations had their session-average peak within ±8 Hz of the cross-site, cross-session average gamma-peak frequency. For gamma-peak based analyses (Fig. 6A, 8A,B), activity was averaged in a window ±8 Hz around the peak determined per stimulus, session and site.

For Dataset 1, after the identification of local maxima as described above, the following was done to account for the larger variability in gamma power and frequency in this stimulus set. The largest two peaks in the spectrum were collected for each site and stimulus. To ensure that the identified peak reliably occurred across days, we ran a one-sided permutation test (alpha = 0.05, n = 1000 permutations of all trials, pooled across all sessions) of the average relative power around the identified peak frequency (±8 Hz around the peak) against activity of ±8 Hz around 190 Hz for each site and stimulus. Note that the p-value was used as a threshold rather than for any inference about the population of peaks across sites and stimuli. The test against high-frequency activity rather than the pre-stimulus baseline period was chosen to identify peaks that are reliably larger than any potential spike leakage effects. For a given stimulus, the largest gamma peaks were typically similar across recording sites (Dataset 1: 91% of sites shared a common peak ±16 Hz, see Figure S1A). We used the identified peak frequency per site and stimulus to group sites with a similar peak and subsequently analyze repetition-related effects in these sites and a frequency band ±16 Hz around the peak. Conclusions were similar when grouping only sites with the peaks within ±8 Hz of the largest peak (87% overlap). 49% of sites with the largest peak exhibited a second peak (defined as a peak within ±16 Hz of most common second largest peak, Figures 2, S2).

### Multi-unit analysis (conversion to dMUA, normalization)

Preprocessing to obtain the MUA envelope, a continuous measure of multi-unit spiking, was described in section Recordings (acquisition, filtering). For the calculation of rate modulations (but not spike-field locking measures), the MUA signal was smoothed with a Gaussian kernel (SD 20 ms). We observed that different recording sites exhibited different levels of noise or background activity. We reasoned that the minimal observable activity in a site would constitute the best available estimate of this noise floor. In order to obtain measures of changes in firing activity, we therefore computed dMUA (for “denoised” MUA). We first computed MUA in 10 ms windows over -1s-1.5 s peri-stimulus onset time, separately for all trials. Subsequently, the minimum MUA value observable per site and day was subtracted from the MUA signal to obtain dMUA.

Based on the dMUA signal, changes in firing rate were then obtained in the form stimulation/baseline, using a common baseline across all trials, where the baseline period was −0.95 to 0 s before stimulus onset.

### Statistics: general procedures

Note that in general, as is common in non-human primate neurophysiology, our sample size (N = 2 to 4 animals per Dataset) only allows an inference on our sample (rather than the population) of macaques. Array recordings allow for an inference over either recording sites or sessions (trials). Given the nature of our questions that compared responses between conditions in different sessions, and the possible interdependence of LFP responses over sites, our units of observation were typically sessions. For certain analyses for natural stimuli, our units of observation were stimuli or stimulus-site combinations, to investigate the dependence of repetition effects on stimulus drive. Permutation tests were conducted using 1000 permutations. For a two-sided test, the minimal p-value obtainable is therefore 0.002. In case of spectrally or time-resolved analyses, the same procedure as described for average gamma-band or dMUA responses was applied to all frequency or time bins, and a multiple comparison correction was applied with an alpha per test of 0.05 and a false discovery rate across the multiple tests of 0.05 (Korn et al., 2004).

Multiple linear regression models of the form

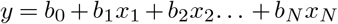

were fit to single-trial gamma-band or MUA responses (y) using N predictors *x*_1_ to *x_N_*. Fits were performed using the Matlab function fitlm. The final model is based on pooled data from all animals, models for individual animals yielded qualitatively similar results. Full models that can include non-significant predictor terms are reported in the text. Dropping these predictors did not qualitatively change any effects. Pairwise correlations between predictors were performed, and in case of predictors with high correlations, only one of the predictors was included in the model. Stimulus identity and recording session number were treated as categorical predictors per animal. Models using random effects (Matlab function fitlme) for animals and sessions yielded qualitatively similar results. Details are discussed below for each dataset.

### Measuring and testing natural image repetition effects on dMUA and gamma responses

We observed that dMUA responses on average showed a rapid decrease for the first few repetitions, followed by a lesser decrease for further repetitions. Gamma-band responses (based on the gamma-peak aligned responses per stimulus, averaged across sites), whose behavior with repetition was not always monotonic, showed an inflection point around repetition 4 on average, from an average decrease to an average increase. To quantify repetition effects, linear slopes were fit to the first 4 (“early”) and the later repetitions (“late”) for each stimulus and animal.

Significance testing was based on non-parametric randomization. Slopes were fit separately to early or late repetitions in each session. In each case, slopes were first fit separately per stimulus and monkey, and the mean value across session was averaged over stimuli and monkeys, providing one observed average slope. This slope was tested against zero based on the distribution of slopes across sessions (and blocks), by randomly flipping the sign of the slope of a given session and stimulus, and computing the mean of these randomized slopes 1000 times. Using the median, rather than the mean, over sessions gave qualitatively the same results. A similar approach was used for statistical testing of the difference in slopes between early and late repetitions. First, the difference between early and late slopes was averaged over stimuli and monkeys. Subsequently, this observed difference was compared to the distribution of differences obtained from a randomization test. In each randomization, a random decision was made to exchange the early versus the late slopes, or not. Subsequently, the slope differences were averaged over stimuli and monkeys, exactly as for the observed data. Across 1000 randomizations, this resulted in a distribution of slope differences against which the observed slope difference was compared.

For Figure 4B, to obtain a metric that normalizes for stimulus-induced changes in power and thereby does not artificially diminish potential effects in lower frequencies, we computed the repetition-related change (RRC) metric. To this end, the linearly fit slopes and intercepts were used to estimate a ratio between the first and the last repetition of a set of trials (early or late): r(last)/(0.5*(r(last)+r(first))). The more direct ratio r(last)/r(first) was avoided since it is less robust when both responses are near zero.

Statistical inferences were based on the procedure described above. In addition, we estimated standard errors of the mean, purely for illustration purposes. The standard error of mean responses was computed using a bootstrap procedure. For 1000 resampling steps, for each stimulus, the responses to a given stimulus repetition (or slope) were resampled, with replacement, from all available sessions. Both the original data and the resampling distribution were then averaged across stimuli, and thereafter across animals. This was done on the cross-site averaged data. One standard error of the mean was then estimated as one standard deviation of the resulting average resampling distribution, all figures display ±2 SEM as either bars or as shaded areas.

### Measuring dependence of natural image repetition effects on stimulus response strength

The degree to which a given recording site changed its response (e.g. dMUA response) to a stimulus may depend on the overall response strength to the stimulus. We therefore analyzed the relation between the repetition-related change and the response strength, separately for early and late repetitions. Two potential pitfalls have to be considered in this analysis. 1) The mean response strength and the repetition-related change can show trivial correlations due to circularity. For example, all other things being equal, a site which shows increasing response strength with repetition will also show a higher mean response. 2) In cases where the response strength is weak (e.g. suppressed below baseline), changes in response strength are potentially limited by a floor effect. To address the first problem, we linearly fit the responses for each session, recording site and stimulus for the repetitions in question (early or late), in a cross-validated manner. The fit

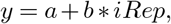

with *iRep* indicating repetition number, yielded an estimate of the intercept *a* at repetition “zero”, as an estimate of response strength without any repetition-related change in response. The fit also yielded aslope *b*. When slope and intercept estimates are based on the same data, another bias can occur due to regression to the mean. We therefore performed two linear fits, each based on a different half of the repetitions (within the early or late categories). One half included every second repetition starting with the first, the other half every second repetition starting with the second, resulting in two independent estimates of slopes and intercepts. Simulations confirmed that this removed the bias. We then tested whether the intercept was predictive of the slope through Spearman’s rank correlation. For each stimulus and recording site combination, we obtained the median slopes and intercepts across sessions. Across the stimulus-site combinations, we then correlated slopes with independently estimated intercepts. We performed this procedure for the two independent combinations of slopes and intercepts and averaged the two resulting correlation values.

To address the second problem, namely a potentially trivial correlation due to a floor effect in stimulus-site combinations that were weakly responsive, we assessed the consistency of the correlation values across a median split of the data by response strength. For each animal, the stimulus-site estimates of the intercept (as our estimate for response strength) were partitioned into two parts with equally many data points (median split), and the correlation was computed for each resulting half of the data. To test for statistical significance, correlations were then averaged for each data half across animals, and the resulting correlation value tested against a permutation distribution (randomizing intercepts 1000 times for each data half per animal, then averaging across animals) using multiple comparison correction across halves (Korn et al., 2004). Note that this procedure is conservative and substantially decreases the absolute value of an existing correlation.

### Measuring stimulus-specific repetition effects for natural images through normalized correlation or linear regression modeling

We reasoned that any stimulus-specific trajectory of a given feature with repetition (e.g. increase, rapid decrease followed by steady response, etc.) would be reflected in correlations between the trajectories across recording sessions (Fig. 5A): the trajectory of stimulus *x*_1_ in session *n*_1_ should correlate more strongly with the trajectory of *x*_1_ in the other sessions than with the trajectory of the other stimuli *x*_2_ to *x_N_* in the other sessions. The trajectory can be computed for arbitrary features, such as LFP power in a specific frequency b and, or MUA responses in a given time bin. By repeating the process for different frequencies or time bins, correlation spectra or time-resolved dMUA correlations were obtained. To compute correlations, we first normalized the trajectory for each session, site and stimulus using a z-score across the repetitions. This results in a trajectory that has a specific shape, but is both demeaned and scaled by the standard deviation of the distribution of repetitions of a stimulus in a session. The z-scored data was then concatenated for each session, using a fixed stimulus order, i.e. irrespective of the actual stimulus order in the session, yielding a “fingerprint” vector of trajectories for each session (see Figure 5A for an illustration of the procedure). The normalization should remove effects that are caused by both greater means and greater variance (that typically accompanies greater mean responses) between different stimuli, in the feature of interest, for example in gamma-band activity. Explorative analyses showed that results based on simple demeaned responses were qualitatively similar to results based on z-scored responses. For a session block to be included in this analysis, it was required that at least 50% of the “fingerprint” vector contained data (this excluded 0-2 session blocks per animal, which were incomplete second blocks of the recording day). Furthermore, for spectral analyses, it was required that at least 5 sites showed a reliable gamma-band peak for a stimulus to be included (this excluded 0 stimuli for LFP gamma power, and 4, 0, or 13 stimuli for MUA-LFP PPC per animal). The data was averaged across sites after normalization. We then used a split-half procedure, where two random halves of the sessions were averaged repeatedly (s = 100 times) and the two average “fingerprint” vectors were correlated. These s split-half Pearson correlations were then averaged to yield a final estimate of the correlation value per animal. Subsequently, values were averaged across animals. To see whether this correlation value was stimulus specific, we used a permutation test. Specifically, for 1000 iterations for each of the s split-halves, we computed pairwise correlations between 1) the intact vector of one session half and 2) the other session half with the trajectories re-ordered in a randomized stimulus order. Using the same procedure as for the observed data, the s split-halves were then averaged. Subsequently, the 1000 permuted values were averaged across animals. The resulting distribution of 1000 correlation values was then used for a two-sided test at alpha = 0.05. For spectra or time-resolved correlations, we used a false discovery rate based multiple comparison correction (FDR = 0.05, alpha = 0.05, Korn et al. (2004)).

#### Multiple regression analysis of repetition effects

The correlation analyses were optimized to remove variance unrelated to stimulus repetition (by z-scoring across repetitions in a repetition sequence per stimulus, session and site, and by then averaging across session halves). Averaging effects across session halves can reduce the effects of various sources of noise, including measurement noise and, given the design, the effects of varying the neighboring a given stimulus, and effects of varying lag between repetitions. We also developed multiple regression models to fit single-trial responses directly, using either dMUA responses or LFP gamma-band peak aligned responses, averaged over sites, in both cases log-transformed so that the fit was not dominated by stimuli with strong responses. Stimulus responses were modeled for each animal individually, and a categorical session (i.e. recording day) regressor was included. Treating these variables as random effects did not substantially affect conclusions.

### Measuring and testing stimulus specificity, location specificity and persistence

Analysis and testing for Datasets 2 and 3 (Figures 6–8, S6-S8) followed a similar logic. Sequences of stimulus repetitions in blocks were generated that kept the overall trial number, stimulus identity, time in session and reward number identical (with the exception of within-session persistence tests). A block of a particular type, e.g. A[B]A or Out[In], constituted a condition. For such a condition, a given neuronal “feature”, such as LFP power relative to baseline in a particular frequency bin or dMUA activity relative to baseline, was first computed for each animal, session, stimulus and site. Activity was then averaged across sites.

To obtain a measure of changes with stimulus repetition, we then fit a linear regression to the feature based on the log-transformed trial number in the block,

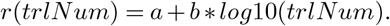

For LFPs, the first 4 trials were excluded for the sake of consistency with the analyses for natural images. From the resulting fit, we computed our measure of “repetition-related change” (RRC, see also previous section “Measuring and testing natural image repetition effects on dMUA and gamma responses”). The same linear regression fit was also used to estimate the “initial response in block”, by using the intercept. The repetition-related change and intercept were averaged first across sessions, then stimuli and animals, thereby giving equal weight to each animal and each stimulus.

Based on fits for each session, stimulus and animal, permutation tests were computed using 1000 permutations, for each frequency bin or time point. To test for the existence of a repetition effect in a given block (i.e. single condition test against zero), a session-level test was performed. The mean of the slopes across sessions was compared against a distribution created by randomly changing the sign of a given session and computing the mean across these randomized slopes 1000 times. Using the median, rather than the mean, over sessions gave qualitatively the same results. The resulting distribution was used to determine two-sided, p<0.01 cut-offs as shown in the bar plots in the insets of Figures 6A, 7A, and observed fits were considered significant, if they fell outside these cut-offs.

For tests regarding the difference between any two blocks (conditions), the difference between conditions was computed on the session level (i.e. separately per animal and stimulus, and for within-session comparisons for each session), and averaged first across sessions, then stimuli and animals. It was tested against a distribution of differences computed the same way, but with condition labels randomly permuted in the first step.

For Dataset 3, fewer sessions were available, and stimuli occurred in a fixed sequence in the session in separate “task blocks”, the purpose of which, together with the breaks between “task blocks”, was to create “mini-sessions”. To obtain a sufficiently large sample for a permutation test, responses were concatenated across “task blocks”, effectively treating them as separate sessions. Subsequently, condition labels were randomly permuted between conditions only for matching stimuli, and averaged across all sessions and stimuli simultaneously. Furthermore, trials were binned in bins of ±3 trials around the trial number in each block.

For illustration purposes, a bootstrap error estimation procedure was used to estimate the variability in the data, for example in Figure 6. To this end, responses were resampled, with replacement, 1000 times from all sessions from the same block and stimulus in each animal, and averaged across the sessions, yielding a resampling distribution for the response in question (e.g. gamma-band response in a trial, LFP repetition-related change in a frequency bin). The resulting distributions were averaged across stimuli and animals. The standard deviation of this distribution is the bootstrap estimate of the standard error of the mean (SEM), and all plots show ±2 SEM. For Dataset 3, data was binned ±3 trials around the trial number in a given block, and bootstrap resampling was performed on these trial bins rather than across sessions, to overcome low session numbers for some animal-block combinations.

Regression modeling for Dataset 2 was performed as follows. The model contained trial-based terms and categorical terms for particular block types. Specifically, the model included a term for the log-transformed number of consecutive repetitions of the same stimulus (“Log(consec. stim. rep. num.”), and terms that indicated if a given block constituted an immediate repetition block (A[A]B, “immediate rep. block”) or delayed repetition block (AB[A], “delayed rep. block”), effectively allowing different offsets to be added to the feature estimate in these cases. Furthermore, interactions (“Log(consec. stim. rep. num.” x “immediate rep. block”), and (“Log(consec. stim. rep. num.” x “delayed rep. block”) were included, effectively allowing a change in the steepness or direction of the repetition effect for these blocks. Note that the value for the main effect of “Log(consec. stim. rep. num.)” will therefore indicate the effect of stimulus repetition, the values for the main effects of “immediate rep. block” and “delayed rep. block” indicate changes in the intercept (i.e. initial block response), and the interaction terms indicate changes in the slope (i.e. difference in the repetition-related change) depending on the block type. In addition, the model contained terms for the animal identity, session identity per animal, stimulus identity per animal, microsaccade rate, pupil response, stimulus duration in the previous trial, inter-stimulus-interval, reaction time, and overall trial number in the session.

Regression modeling for Dataset 3 was performed as follows. To avoid higher-order interactions due to the trivial effect of stimulus location, and to decorrelate the location-specific from the total trial number in a task block as much as possible, only trials where the RF was stimulated were modeled, and only for the blocks that dissociated location-specific or persistence effects directly (Out[In] vs In[In] and OutOut[In] vs InOut[In] respectively, corresponding to the contrasts in Figure 8). This way, a significant contribution of the “Log(local rep. num.)” regressor dissociates location-specific from general repetition effects (tested using “Log(task block trial num.)” for blocks Out[In] compared to In[In], whereas a significant contribution of the “Log(local rep. num.)” regressor predicts an increase in gamma power for block InOut[In] compared to OutOut[In], indicative of persistence. In addition, the model contained terms for the animal identity, session identity per animal, stimulus identity per animal, microsaccade rate, pupil response, stimulus duration in the previous trial, inter-stimulus-interval, reaction time, and overall trial number in the session.

