## Supplementary material for "Stimulus-specific plasticity of macaque V1 spike rates and gamma": Suppl. Figures

November 13, 2020

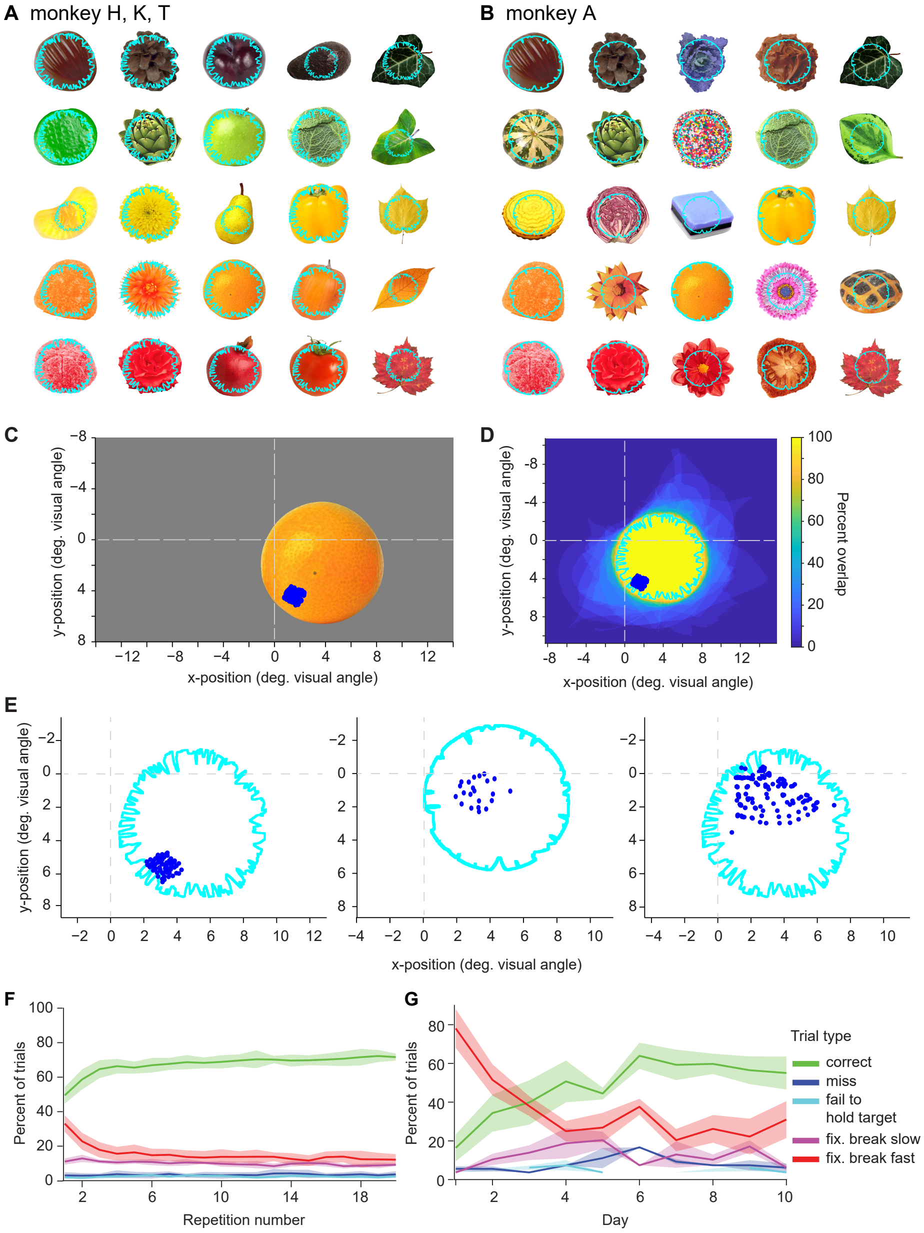

Figure S1: (A) Stimuli used for monkeys H, K and T. Stimuli were selected from a large library (Hemera Photo-Objects, Hemera Technologies). (B) Stimuli for monkey A. Cyan outlines on each stimulus illustrate the overlap with all other images used for a given animal, and thereby the area where a change could occur. Note that stimuli are individually scaled for this figure. (C,D) Illustration of stimulus position with respect to fixation (i.e. 0,0 refers to the fixation point) and the receptive field centers of the V1 array for an example animal (monkey H). (C) Stimulus position on the monitor for an example stimulus. (D) Illustration of percentage of overlap between all images, the 100% region is indicated with cyan outlines. (E) Illustration of overlap of all stimuli with respect to fixation and the receptive field centers of the V1 recording sites for monkeys K, A, T (from left to right). (F,G) Several parameters of behavioral performance as a function of repetition number (F) and as a function of recording day (G). Behavioral performance parameters were: Percentage of all trials that were hits, misses, failures to hold target (i.e. trials during which the animal first saccaded to the change location but did not fixate it for long enough), fixation breaks that were slow ($>$1 s after stimulus onset) or fast ($<$1s after stimulus onset). Error regions show $\pm$ 1 SEM across animals. Animals were more likely to respond with rapid fixation breaks during the first few presentations of a novel stimulus, especially on the first few days. Note that due to the design of the experiment, session-novel stimuli were introduced over the entire course of the session, so this behavior cannot be explained by increased fixation breaks at the beginning of the session. Multiple regression modeling of correct versus incorrect trials confirmed that stimulus repetition number, but not total trial number, affected whether a correct trial would be performed, and this held only when fixation break trials were included in the data ($p<4.94e-56$ for stim. rep. num., $p=0.26$ for total trial number). Reaction times showed no significant relationship to stimulus repetition, nor to total trial number (all $p>0.3$).

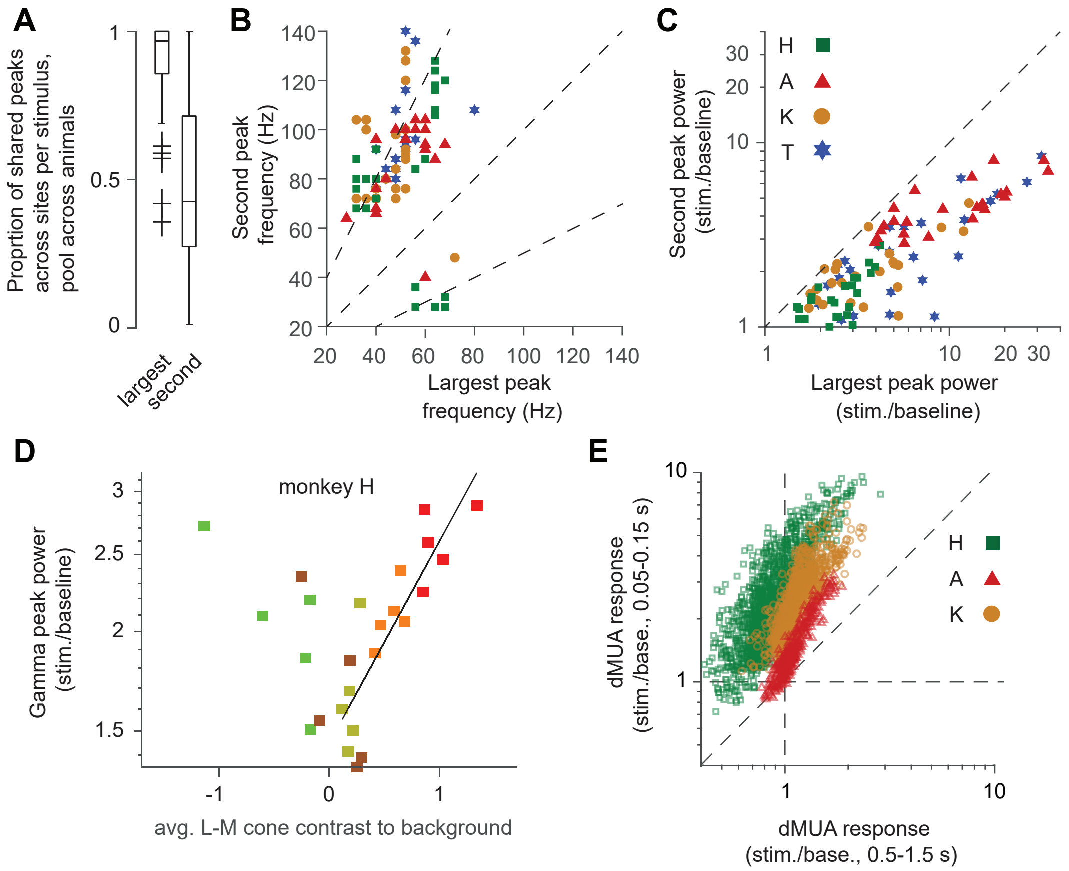

Figure S2: (A) Prevalence of LFP gamma-band peaks shared between sites (within $\pm$ 16 Hz, see Methods) for a stimulus, across stimuli and animals. Largest peak refers to the largest peak most common across recording sites. Second peak refers to the most common second largest peak among recording sites exhibiting the largest peak. Horizontal lines refer to median, 25th and 75th percentile. (B) Peak frequency of the largest peak versus second largest peak. Each symbol indicates average gamma-band peak frequency for a stimulus and animal. Dashed lines indicate the ratios 1:2, 1:1 and 2:1. There are clusters around peak frequency ratios of 2:1 and 1:2. These precise frequency ratios would be consistent with harmonic relationships, yet note substantial scatter around the respective lines (see also Methods). (C) Fold change in power at the gamma-band peak and second largest peak for all stimuli. (D) Dependence of relative gamma-band power on L-M cone contrast for an example animal (monkey H). Each symbol shows average gamma-band power around the largest peak per stimulus across sessions and sites. Color of the symbol is an approximation of the stimulus color (yellow stimuli are shown in a darker hue) to give an intuition of the color dependence. Correlation (Pearson's r) between positive L-M cone contrast and gamma power was $r = 0.88$ for this animal ($r = 0.85$ on average across monkeys, range 0.72-0.93, $p<0.002$). (E) Average dMUA fold change responses (stimulation/baseline) during the initial transient and responses during the post-transient period for each recording site-stimulus combination.

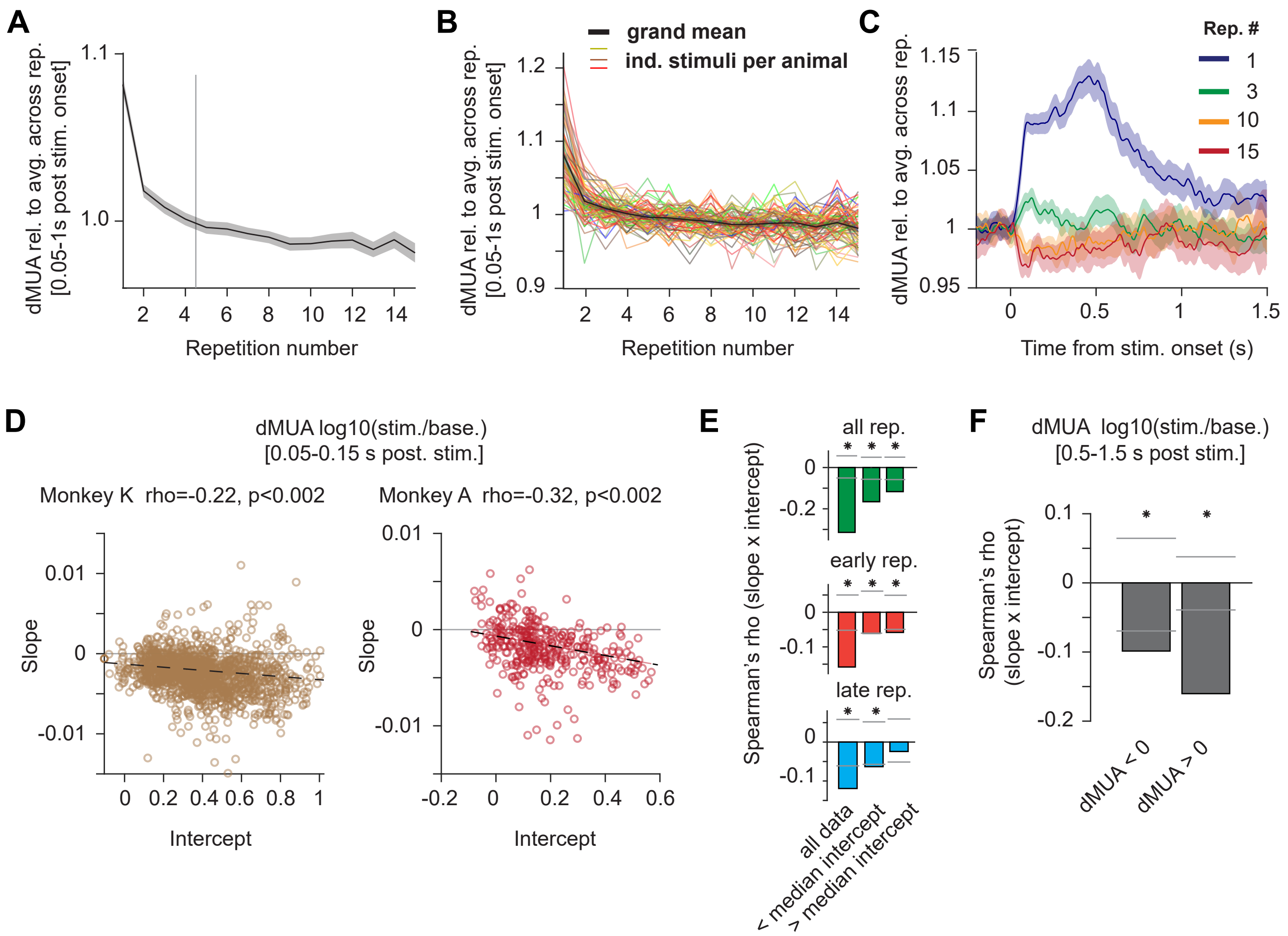

Figure S3: (A) Firing rate responses computed as in 3B for the black line, but normalized by dividing by the mean across repetitions. The vertical line indicates cutoff between "early" and "late" repetitions. (B) Same as black line in (A), with separate lines for each stimulus and animal (line color was chosen to approximate the dominant color in the stimulus as in 3C). (C) Grand average time-resolved dMUA responses as in 3D, but normalized as (A, B), for repetitions 1, 3, 10 and 15. (D) Same as Figure 3F, but for monkey K and monkey A, respectively. (E) Spearman correlation coefficients between slopes and intercepts, across recording sites and stimuli, averaged over animals, separately for all repetitions (top panel), early repetitions (middle panel) and late repetitions (bottom panel), and based on all stimulus-site combinations (first bar), or only the data below (second bar) or above (third bar) the median intercept. (F) Same as the first bar in the top panel of (E), but for the late time window (0.5- 1.5 s post-stimulus), and separately for intercepts below or above baseline (- 0.5-0 s pre-stimulus onset) activity. (A-C) Shaded error regions indicate a bootstrap estimates of $\pm 2$ SEM across sessions (see Methods). (E,F): Gray horizontal lines indicate two-sided significance thresholds at p=0.01, based on a permutation test. Stars indicate significance ($p<0.05$, two-sided, corrected for multiple comparisons.)

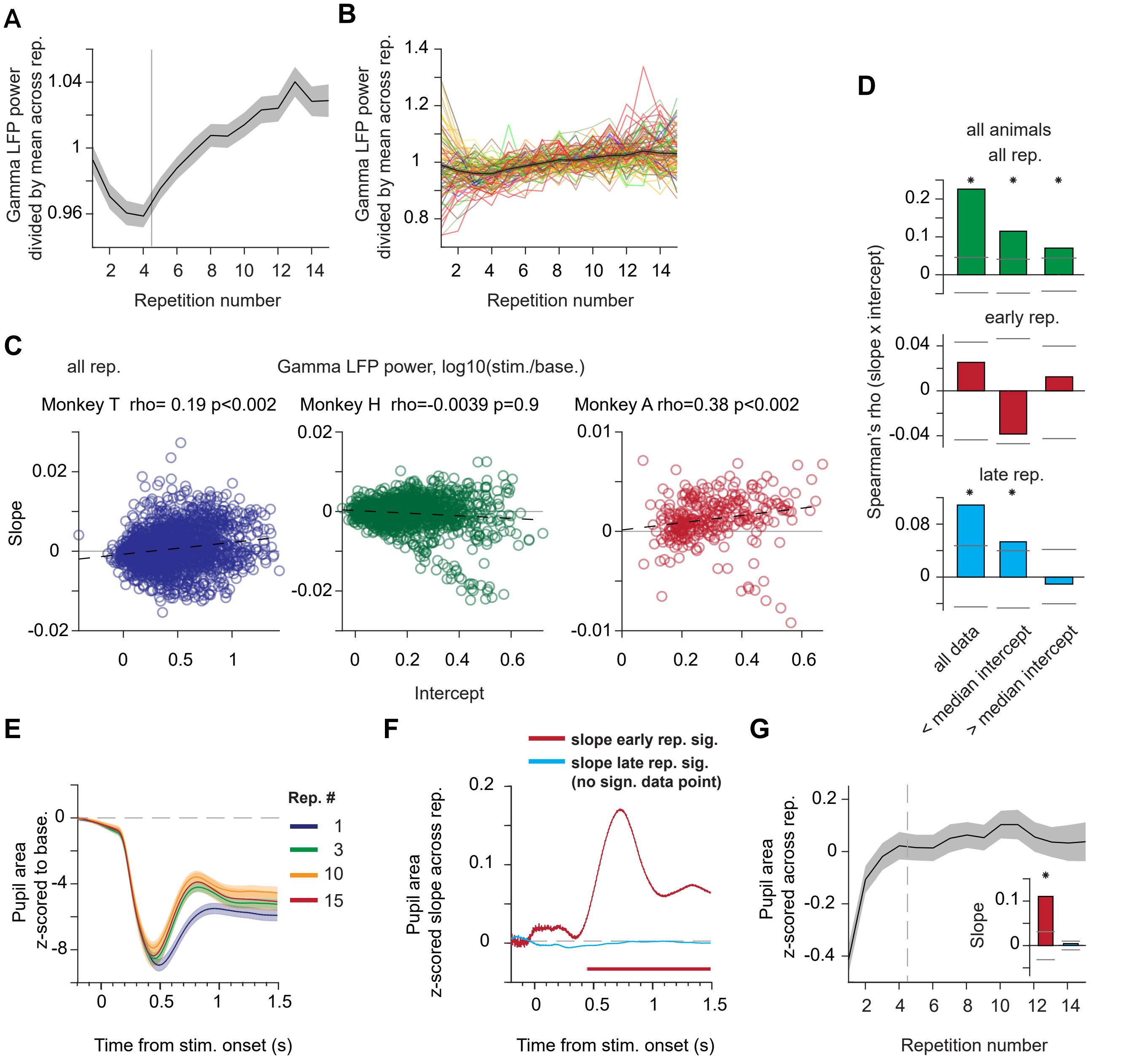

Figure S4: (A) Same as Figure 4, but normalized by dividing by the mean across repetitions. The vertical line indicates cutoff between "early" and "late" repetitions. (B) Same as black line in (A), with separate lines for each stimulus and animal (line color was chosen to approximate the dominant color in the stimulus). (C) Same as Figure 4E, but for monkey T, monkey H, and monkey A, respectively. (D) Spearman correlation coefficients between slopes and intercepts, across recording sites and stimuli, averaged over animals, separately for all repetitions (top panel), early repetitions (middle panel) and late repetitions (bottom panel), and based on all stimulus-site combinations (first bar), or only the data below (second bar) or above (third bar) the median intercept. (E) Pupil area as a function of time from stimulus onset, separately for repetitions 1, 3, 10 and 15 (see Methods). (F) Slopes of linear fits to pupil responses of early (red) or late (cyan) repetitions, computed as in Figure 3E. Colored bar indicates multiple-comparison corrected, significantly positive slopes (i.e. decreasing amount of pupil constriction) of linear fits to early repetitions. There was no significant effect for late repetitions. (G) Pupil area (0.5-1.5 s from stimulus onset) as a function of stimulus repetition number, z-scored as in (F). (A,B,E,G) Shaded error regions indicate bootstrap estimates of $\pm$ 2 SEM across sessions (see Methods). (D) Gray horizontal lines indicate two-sided significance thresholds at p=0.01 based on a permutation test. Stars indicate significance ($p<0.05$, two-sided, corrected for multiple comparisons).

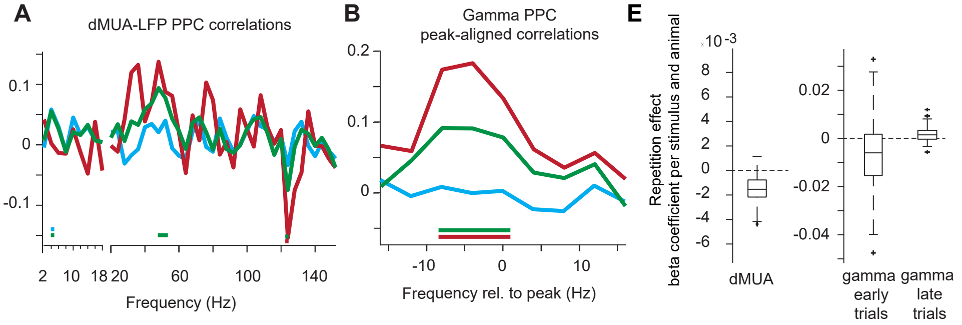

**C**

**Regression model of dMUA response (all trials)**

| | SumSq | DF | MeanSq | F | pValue | expVar | expVarPart | beta coefficient |
|---|---|---|---|---|---|---|---|---|
| Stimulus | 41.68 | 72 | 0.58 | 737.09 | <1E-65 | 0.7724 | 0.9221 | |
| Session | 1.40 | 44 | 0.03 | 40.56 | <1E-65 | 0.0260 | 0.0310 | |
| Lag | 0.03 | 1 | 0.03 | 42.21 | 8.55E-11 | 0.0006 | 0.0007 | -0.0008 |
| Lag:stimulus | 0.16 | 74 | 0.00 | 2.80 | 2.02E-14 | 0.0030 | 0.0036 | |
| Log(Overall trial number) | 0.06 | 1 | 0.06 | 80.85 | 2.82E-19 | 0.0012 | 0.0014 | 0.0053 |
| Log(Overall trial number):stimulus | 1.12 | 74 | 0.02 | 19.22 | <1E-65 | 0.0207 | 0.0247 | |
| Log(Repetition num.) | 0.53 | 1 | 0.53 | 672.60 | <1E-65 | 0.0098 | 0.0117 | -0.0008 |
| Log(Repetition num.): stimulus | 0.21 | 74 | 0.00 | 3.68 | 4.29E-24 | 0.0040 | 0.0047 | |
| unexplained | 8.76 | 11160 | | | | | | |

**D**

**Regression model of gamma power (early trials)**

| | SumSq | DF | MeanSq | F | pValue | expVar | expVarPart | beta coefficient |
|---|---|---|---|---|---|---|---|---|
| Stimulus | 114.93 | 96 | 1.20 | 244.31 | <1E-65 | 0.7723 | 0.9012 | |
| Session | 7.18 | 62 | 0.12 | 23.63 | <1E-65 | 0.0482 | 0.0563 | |
| Lag | 0.23 | 1 | 0.23 | 46.25 | 1.18E-11 | 0.0015 | 0.0018 | 0.0056 |
| Lag:stimulus | 0.52 | 99 | 0.01 | 1.08 | 2.78E-01 | 0.0035 | 0.0041 | |
| Log(Overall trial number) | 1.03 | 1 | 1.03 | 210.62 | 1.22E-46 | 0.0069 | 0.0081 | 0.0525 |
| Log(Overall trial number):stimulus | 2.31 | 99 | 0.02 | 4.75 | 1.70E-46 | 0.0155 | 0.0181 | |
| Repetition num. | 0.29 | 1 | 0.29 | 58.87 | 2.07E-14 | 0.0019 | 0.0023 | -0.0206 |
| Repetition num: stimulus | 1.05 | 99 | 0.01 | 2.16 | 4.75E-10 | 0.0070 | 0.0082 | |
| unexplained | 21.29 | 4344 | | | | | | |

**Regression model of gamma power (late trials)**

| | SumSq | DF | MeanSq | F | pValue | expVar | expVarPart | beta coefficient |
|---|---|---|---|---|---|---|---|---|
| Stimulus | 317.49 | 96 | 3.31 | 761.64 | <1E-65 | 0.80089 | 0.91413 | |
| Session | 20.24 | 61 | 0.33 | 76.42 | <1E-65 | 0.05106 | 0.05828 | |
| Lag | 1.39 | 1 | 1.39 | 319.41 | <1E-65 | 0.00350 | 0.00399 | 0.0087 |
| Lag:stimulus | 1.71 | 99 | 0.02 | 3.97 | 8.526E-36 | 0.00430 | 0.00491 | |
| Log(Overall trial number) | 1.60 | 1 | 1.60 | 368.68 | <1E-65 | 0.00404 | 0.00461 | 0.0130 |
| Log(Overall trial number):stimulus | 3.88 | 99 | 0.04 | 9.03 | <1E-65 | 0.00979 | 0.01117 | |
| Repetition num. | 0.23 | 1 | 0.23 | 52.72 | 4.09E-13 | 0.00058 | 0.00066 | -0.0003 |
| Repetition num: stimulus | 0.78 | 99 | 0.01 | 1.81 | 1.652E-06 | 0.00196 | 0.00224 | |
| unexplained | 49.11 | 11310 | | | | | | |

Figure S5: (A) Correlation spectra for dMUA-LFP PPC. (B) Gamma-peak aligned versions of (A). (C-E) Multiple linear regression models to test stimulus specificity in natural image paradigm (see Methods for modeling approach). Stimulus specificity is shown by a significant interaction between repetition number and stimulus identity (Repetition num: stimulus). Note that the direction of the main effect of repetition is not directly interpretable due to the interaction term. The direction of the net effect (main effect plus interaction) is shown in (E), as box-and-whisker plots across stimuli. (C) Model for dMUA responses of the initial transient response period (0.05-0.15 s post stimulus onset, log-transformed) based on all 15 repetitions. Qualitatively similar results were obtained for analyses averaging across the entire stimulus time period, or across the time period for which gamma-band responses were computed (0.5-1.5 s post stimulus onset). There was a significant effect of the log-transformed stimulus repetition number, which showed significant modulation by stimulus identity (significant interaction term). Net effects were negative (see (E)). The present model based on log-transformed repetition numbers and all repetitions was chosen to capture the repetition effect in dMUA responses in a single model. The qualitative pattern of results, and the significant effect of repetition number, remained when analyses were performed for individual animals, for the non-log-transformed repetition number, when including the pupil response for animals where it was available, and when models were fit only to the initial 4 or only to the late repetitions (log- transformed or not). Models fit to only the late repetitions still showed significant effects of repetition. (D) Models for gamma-band responses (log-transformed, see Methods). In contrast to the dMUA responses, and in line with the previous analyses on the gamma-band, models fit to the early compared to the late repetitions showed a qualitative change in the repetition effect, such that two models will be presented here. Both models showed significant effects of stimulus repetition. In the model for early repetitions (upper table), net repetition effects for individual stimuli were predominantly negative. In contrast, net effects were smaller and predominantly positive for late repetitions (see (E)). For both models, there was a significant effect of the log of the total trial number in the session, which tended to be positive (as compared to negative or absent in the paradigms using gratings). Both the qualitative pattern of results and the significant effect of repetition number were unchanged when analyses were performed for individual animals, and when including the pupil response for animals where it was available. SumSq = sum of square of variation, DF = degrees of freedom, MeanSq= SumSq average per DF used, F = F-value of ANOVA, pValue = significance value, expVar = explained variance of a regressor, expVarPart = share of explained variance of a regressor normalized by entire explained variance, beta coefficient = value of regressor in multiple regression model. Non-significant regressors are shown in gray. (E) Distribution of net repetition effects (i.e. main effect plus individual effect per stimulus and animal) across stimuli for dMUA from the model in (C) and for gamma from models in (D).

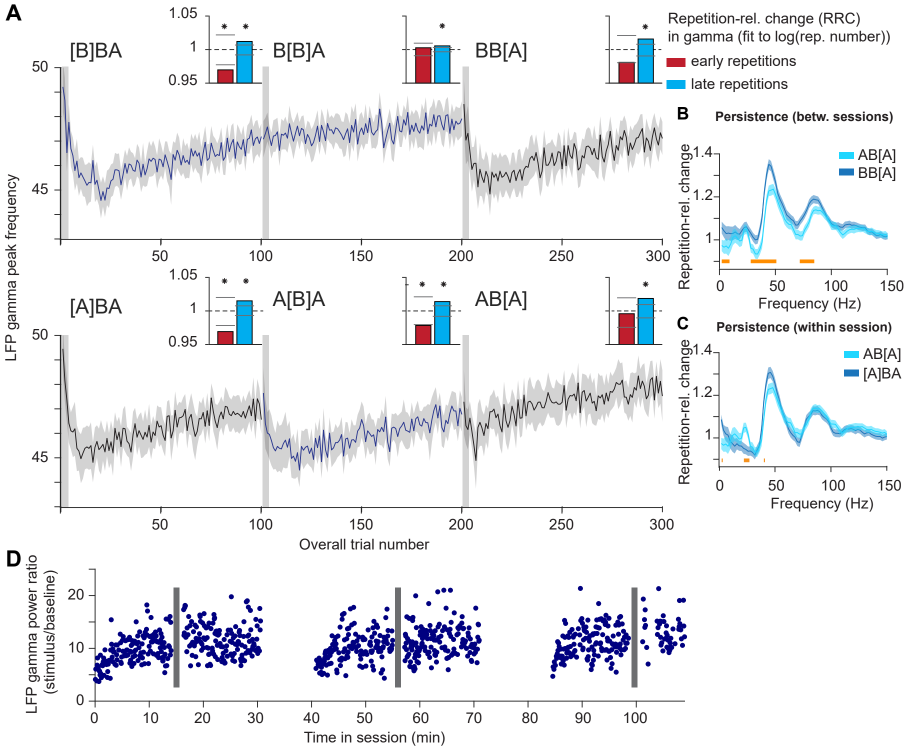

**E**

| | SumSq | DF | MeanSq | F | pValue | expVar | expVarPart | beta coefficient |
|---|---|---|---|---|---|---|---|---|
| stimulus | 47021.80 | 2 | 23510.90 | 2171.04 | <1e-65 | 0.101 | 0.323 | |
| stimulus:monkey | 34174.72 | 2 | 17087.36 | 1577.88 | <1e-65 | 0.073 | 0.235 | |
| monkey:sessInd | 27084.74 | 71 | 381.48 | 35.23 | <1e-65 | 0.058 | 0.186 | |
| stim. duration prev. trial | 30.06 | 1 | 30.06 | 2.78 | 9.57E-02 | 0.000 | 0.000 | -0.14 |
| inter-stimulus-interval | 341.67 | 1 | 341.67 | 31.55 | 1.96E-08 | 0.001 | 0.002 | -0.18 |
| pupil response | 2.45 | 1 | 2.45 | 0.23 | 6.34E-01 | 0.000 | 0.000 | 0.00 |
| microsaccade rate | 122.84 | 1 | 122.84 | 11.34 | 7.58E-04 | 0.000 | 0.001 | -0.12 |
| Log(consec. Stim. Rep. num.) | 30935.14 | 1 | 30935.14 | 2856.61 | <1e-65 | 0.066 | 0.213 | 3.39 |
| Log(Overall trial num.) | 638.56 | 1 | 638.56 | 58.97 | 1.65E-14 | 0.001 | 0.004 | -0.47 |
| immediate rep. block | 2953.78 | 1 | 2953.78 | 272.76 | 5.32E-61 | 0.006 | 0.020 | 11.28 |
| delayed rep. block | 79.13 | 1 | 79.13 | 7.31 | 6.87E-03 | 0.000 | 0.001 | 2.60 |
| Log(consec.stim.rep. num.) X imm. rep. block | 958.13 | 1 | 958.13 | 88.48 | 5.50E-21 | 0.002 | 0.007 | -5.71 |
| Log(consec.stim.rep. num.) X del. rep. block | 1104.57 | 1 | 1104.57 | 102.00 | 6.07E-24 | 0.002 | 0.008 | -1.49 |

Figure S6: (A) LFP gamma peak frequency (see Methods) as a function of overall trial number. Similar to Figure 6A, but for gamma peak frequency instead of gamma power. Insets show repetition-related gamma-frequency changes for early repetitions (red) and late repetitions (cyan) (bars show permutation-based two-sided significance thresholds for p<0.01). There was a peak frequency decrease for early stimulus presentations in most cases, and a peak frequency increase for late repetitions. (B) Repetition-related change in LFP power for blocks AB[A] versus BB[A], i.e. using the between-sessions comparison. (C) Same as (B) but for blocks AB[A] versus [A]BA, i.e. using the within-session comparison. (D) The effect of interleaving many different stimuli and the effect of a rest from the task was tested in a single session in monkey H. A vertical moving grating was repeatedly presented, and after 150 presentations, there was a period of 125 rapid, 200 ms long presentations of grating stimuli with other orientations (denoted by gray bars, see Methods for details), after which stimulation with the original grating continued. This experiment was repeated two additional times in the same session, with periods of 10 min of rest in between stimulation blocks. The time course of gamma across repetitions suggests 1) persistence over intervening stimulation and 2) reset after 10 min of rest. (E) Multiple linear regression model evaluated with ANOVA (see Methods for modeling approach) confirms effects of stimulus repetition and persistence for gamma-band power based on single-trial analyses. The model showed (all p<0.01): 1) a main effect of stimulus repetition, i.e. an increase in gamma-band response with the log-transformed repetition number ("Log(consec. stim. rep. num.)"); 2) stimulus specificity, i.e. an increase in the initial response in the block for the immediate repetition block ("immediate rep. block"), and also a net decrease in gamma-band responses for this block for further stimulus repetitions ("Log(consec. stim. rep. num.)x imm. rep. block"); 3) a persistence effect, i.e. an increase in the initial response for block AB[A], and a reduced increase in gamma for the following repetitions ("delayed rep. block" and interaction). The model controlled for the effects of overall trial number, pupil responses, microsaccade rates, inter-stimulus-intervals, stimulus duration in the previous trial, as well as the stimulus, session and monkey identity (the latter subsume many beta coefficients and capture a lot of variance due to overall differences in response strength to a stimulus for an animal and session). SumSq = sum of square of variation, DF = degrees of freedom, MeanSq= SumSq average per DF used, F = F-value of ANOVA, pValue = significance value, expVar = explained variance of a regressor, expVarPart = share of explained variance of a regressor normalized by entire explained variance, beta coefficient = value of regressor in multiple regression model. Non-significant regressors are shown in gray.

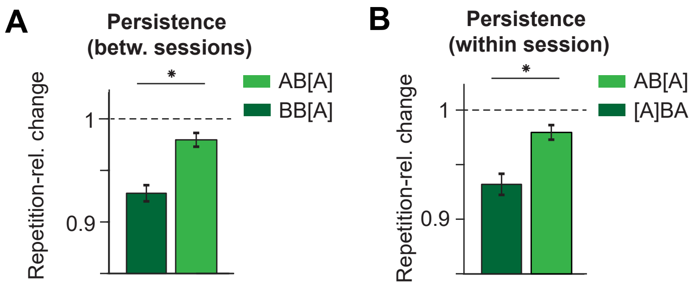

**C**

| | SumSq | DF | MeanSq | F | pValue | expVar | expVarPart | beta coefficient |
|---|---|---|---|---|---|---|---|---|
| stimulus | 31.79 | 2 | 15.90 | 252.99 | <1E-65 | 0.010 | 0.015 | |
| stimulus:monkey | 147.72 | 2 | 73.86 | 1175.50 | <1E-65 | 0.045 | 0.069 | |
| monkey:sessInd | 1870.34 | 72 | 25.98 | 413.42 | <1E-65 | 0.572 | 0.874 | |
| stim. duration prev. trial | 1.42 | 1 | 1.42 | 22.67 | 1.94E-06 | 0.000 | 0.001 | -0.04 |
| inter-stimulus-interval | 4.06 | 1 | 4.06 | 64.68 | 9.38E-16 | 0.001 | 0.002 | 0.02 |
| pupil response | 0.00 | 1 | 0.00 | 0.02 | 8.98E-01 | 0.000 | 0.000 | 0.00 |
| microsaccade rate | 11.90 | 1 | 11.90 | 189.36 | 7.25E-43 | 0.004 | 0.006 | 0.02 |
| Log(consec. Stim. Rep. num.) | 61.03 | 1 | 61.03 | 971.27 | <1E-65 | 0.019 | 0.029 | -0.18 |
| Log(Overall trial num.) | 2.56 | 1 | 2.56 | 40.80 | 1.72E-10 | 0.001 | 0.001 | -0.04 |
| immediate rep. block | 2.25 | 1 | 2.25 | 35.77 | 2.26E-09 | 0.001 | 0.001 | -0.24 |
| delayed rep. block | 6.59 | 1 | 6.59 | 104.89 | 1.51E-24 | 0.002 | 0.003 | -0.11 |
| Log(consec.stim.rep. num.) X imm. rep. block | 0.30 | 1 | 0.30 | 4.85 | 2.77E-02 | 0.000 | 0.000 | 0.13 |
| Log(consec.stim.rep. num.) X del. rep. block | 0.15 | 1 | 0.15 | 2.42 | 1.19E-01 | 0.000 | 0.000 | 0.02 |
| unexplained | 1128.68 | 17963 | | | | | | |

Figure S7: (A) Test for persistence. Repetition-related changes as in Figure 7C, but for blocks AB[A] versus BB[A]. (B) Same as (A) but for blocks AB[A] versus [A]BA. (C) Multiple regression model (see Methods for modeling approach) confirms effects of stimulus repetition and of persistence for dMUA during the stimulus transient based on single-trial analyses. See legend of Figure S6 for further explanation of model and regressor names.

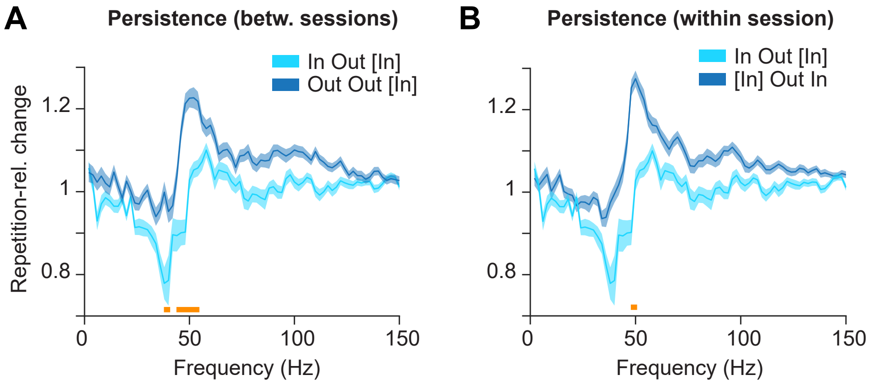

**C** **Regression model of blocks In[In] and Out[In] as in Figure 8A, testing location specificity**

| | SumSq | DF | MeanSq | F | pValue | expVar | expVarPart | beta coefficient |
|---|---|---|---|---|---|---|---|---|
| stimulus | 1038.96 | 2 | 519.48 | 113.27 | 4.46E-48 | 0.062 | 0.302 | |
| stimID:monkey | 1539.03 | 2 | 769.52 | 167.78 | 1.20E-69 | 0.093 | 0.448 | |
| monkey:sessInd | 450.90 | 1 | 450.90 | 98.31 | 8.25E-23 | 0.027 | 0.131 | |
| stimulus duration | 1.43 | 1 | 1.43 | 0.31 | 5.76E-01 | 0.000 | 0.000 | 0.061 |
| inter-stimulus-interval | 19.75 | 1 | 19.75 | 4.31 | 3.81E-02 | 0.001 | 0.006 | -0.043 |
| pupil response | 2.15 | 1 | 2.15 | 0.47 | 4.93E-01 | 0.000 | 0.001 | 0.003 |
| microsaccade rate | 6.48 | 1 | 6.48 | 1.41 | 2.35E-01 | 0.000 | 0.002 | -0.273 |
| Log(Task block trial num.) | 0.94 | 1 | 0.94 | 0.20 | 6.51E-01 | 0.000 | 0.000 | -0.531 |
| Log(Local rep. num.) | 377.24 | 1 | 377.24 | 82.25 | 2.16E-19 | 0.023 | 0.110 | 0.566 |
| Log(overall trial num.) | 0.00 | 1 | 0.00 | 0.00 | 9.82E-01 | 0.000 | 0.000 | -0.670 |
| unexplained | 13194.89 | 2877 | | | | | | |

**D** **Regression model of blocks InOut[In] and OutOut[In] as in Figure 8B, testing persistence**

| | SumSq | DF | MeanSq | F | pValue | expVar | expVarPart | beta coefficient |
|---|---|---|---|---|---|---|---|---|
| stimulus | 717.30 | 2 | 358.65 | 55.59 | 5.11E-24 | 0.062 | 0.391 | |
| stimID:monkey | 830.06 | 2 | 415.03 | 64.33 | 1.56E-27 | 0.072 | 0.452 | |
| monkey:sessInd | 152.93 | 1 | 152.93 | 23.70 | 1.24E-06 | 0.013 | 0.083 | |
| stimulus duration | 0.29 | 1 | 0.29 | 0.05 | 8.32E-01 | 0.000 | 0.000 | -0.096 |
| inter-stimulus-interval | 24.31 | 1 | 24.31 | 3.77 | 5.24E-02 | 0.002 | 0.013 | -0.027 |
| pupil response | 13.34 | 1 | 13.34 | 2.07 | 1.51E-01 | 0.001 | 0.007 | 0.001 |
| microsaccade rate | 4.39 | 1 | 4.39 | 0.68 | 4.10E-01 | 0.000 | 0.002 | -0.282 |
| Log(Task block trial num.) | 0.73 | 1 | 0.73 | 0.11 | 7.37E-01 | 0.000 | 0.000 | 0.225 |
| Log(Local rep. num.) | 50.76 | 1 | 50.76 | 7.87 | 5.10E-03 | 0.004 | 0.028 | 1.115 |
| Log(overall trial num.) | 41.99 | 1 | 41.99 | 6.51 | 1.08E-02 | 0.004 | 0.023 | -0.003 |
| unexplained | 9710.12 | 1505 | | | | | | |

Figure S8: (A) Repetition-related change spectrum for late repetitions, for InOut[In] versus OutOut[In]. (B) Same as (A) but for InOut[In] versus [In]OutIn. (C) Regression model testing location specificity using blocks Out[In] and In[In] (see Methods). The location-specific presentation number, but not the total presentation number in the task block was predictive of gamma power. (D) Regression model testing persistence using blocks InOut[In] and OutOut[In] (see Methods). The location-specific presentation number, which discriminates the block types and predicts more gamma-power for block InOut[In] than OutOut[In], i.e. persistence, was significant.
